# The structure, folding kinetics and dynamics of long poly(UG) RNA

**DOI:** 10.1101/2025.04.22.650042

**Authors:** Riley J. Petersen, Rahul Vivek, Marco Tonelli, Saeed Roschdi, Samuel E. Butcher

## Abstract

Long poly(UG) or “pUG” dinucleotide repeats are abundant in eukaryotic transcriptomes. Thousands of human genes have pUGs longer than 24 repeats, including the cancer-associated lncRNA NEAT1. In *C. elegans*, enzymatic addition of long pUGs to RNA 3′ ends (pUG tails) marks RNAs as vectors of gene silencing. Gene silencing requires at least one pUG fold, a left-handed quadruplex structure that incorporates 12 repeats, but longer pUG tails are more active. Here, we investigate the structure, folding kinetics and dynamics of long pUG RNAs. RNAs with 24 or more repeats slowly form compact, double pUG folds. The forward rate of pUG fold formation *in vitro* is length-dependent with a half-life (t_1/2_) of 13 minutes, while the unfolding rate is very slow (t_1/2_ ∼5 days). Long pUG RNAs display biphasic dynamics with an additional, faster unfolding phase (t_1/2_ ∼30 min). The amplitude of the faster phase indicates partial unfolding. From these data we propose a dynamic model for segmental register exchange and double pUG fold formation in long pUG RNAs. These data broaden our understanding of the structure and dynamics of long pUG RNAs and have implications for understanding the roles of pUG folds in biology and disease.

## Introduction

Poly(UG) or pUG RNAs are dinucleotide simple sequence repeats (SSRs) that are abundant in eukaryotes. SSRs make up ∼3% of the human genome (1) and the most common SSR in mammals is GT/AC (2). Polymorphisms in GT/AC SSRs have been linked to diseases including cystic fibrosis (3,4), asthma (5), cardiovascular disease (6), schizophrenia (7,8) and colorectal cancer (9). GT dinucleotide repeats are more often transcribed than AC repeats as they tend to be downstream of promoter regions (10). Thus, eukaryotic transcriptomes are enriched in GU repeats. These repeats can begin with either a G or U and are collectively referred to as poly(UG) or “pUG” RNAs (11). Previously, we discovered the pUG fold, an unusual left-handed quadruplex (G4) that incorporates 12 UG repeats (Figure 1)(11-13). Humans have ∼20,000 pUGs with 12 or more repeats (11). Among all dinucleotide SSRs with 12 or more repeats in humans, pUGs are the only ones that have a non-random distribution and are enriched near 5′ and 3′ splice sites (11). Expansion of a pUG repeat in the CFTR gene causes mis-splicing and a rare hereditary form of cystic fibrosis with male infertility (3,14). The long noncoding RNA (lncRNA) nuclear paraspeckle assembly transcript 1 (NEAT1) has 29.5 pUG repeats and is overexpressed in many cancers (15-19).

**Figure 1.**
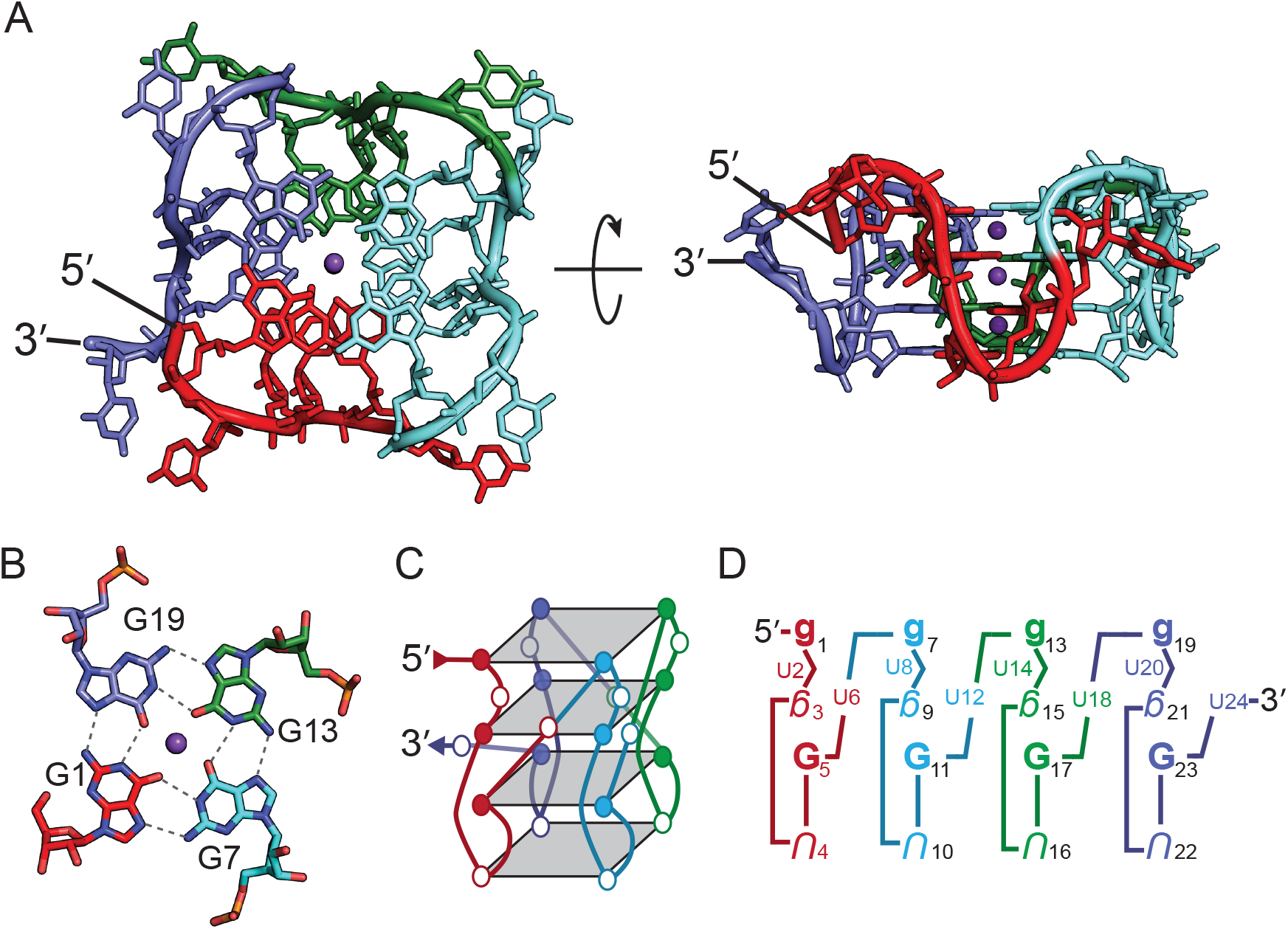
(A) Structure of the (GU)12 pUG fold (PDB: 8TNS, 7MKT). The left-handed G4 is comprised of four segments, each with 6 GU repeats. The segments are color coded from 5′ to 3′ as red, cyan, green and blue. Three central potassium ions are shown in purple. (B) The G1-quartet. (C) Schematic of the pUG fold topology, with G and U nucleotides shown as filled and open circles respectively. (D) Short-hand description of the pUG-fold. Upper case nucleotides are *Anti*, lower case are *Syn*. Bold nucleotides are 2′-endo and italicized nucleotides are 3′-endo. Inverted nucleotides indicate inverted strand polarity relative to the 5′ nucleotide. Small Us represent bulged or looped nucleotides.

The SSR distribution in *C. elegans* is unusual, as pUGs are nearly absent within the encoded transcriptome but instead are post-transcriptionally added to RNA 3′ ends by the enzyme RDE-3 (20,21). These “pUG tails” mark RNAs as vectors for gene silencing and can be over 100 nucleotides long (20,21). Gene silencing in *C. elegans* requires the pUG fold (Figure 1) (11). The unique left-handed G4 topology of the pUG fold recruits the RNA dependent RNA polymerase (RdRP) RRF-1 to RNA 3′ ends for secondary siRNA synthesis (11,20). These siRNAs can target additional mRNAs for cleavage and “pUGylation” in a feed-forward amplification cycle (20). The pUG-tailed mRNAs are packaged in embryos and inherited. A single injection of a pUG tailed RNA into an adult animal promotes gene silencing for 6 generations, a phenomenon known as transgenerational epigenetic inheritance (TEI) (20). The pUG-mediated siRNA amplification pathway provides an explanation as to why GT repeats are very rare in the *C. elegans* genome (11).

The pUG fold has 3 G quartets, 1 U quartet, single uridine bulges and single uridine loops (11-13) (Figure 1). The pUG fold forms around a core of potassium ions and is the only known left-handed RNA G4, owing to a Z-form like backbone configuration in the center of the fold (12). The pUG fold can tolerate insertion of AA dinucleotides while maintaining its structure and gene silencing activity (11). Intramolecular pUG folds are not restricted to RNA 3′ ends and can form in the middle of RNAs *in vitro* (11). The pUG fold requires 12 guanosines (11). Active pUG tails also minimally require 12 guanosines, but gene silencing activity increases with pUG tail length and plateaus somewhere between 18.5 and 40.5 repeats (11).

In this study we examine the structure, folding kinetics and dynamics of pUG RNAs as a function of length. Circular dichroism (CD) spectroscopy, nuclease mapping, native gel electrophoresis, nuclear magnetic resonance (NMR) spectroscopy and molecular dynamics (MD) simulation experiments show that long pUGs with ≥ 24 repeats can form compact double pUG folds. *In vitro* folding of long pUG RNAs results in a stochastic mixture of single and double pUG folds that slowly resolve into double pUG folds over the course of hours. The forward folding rate of the pUG fold depends on the number of repeats and has a half-life measured in minutes. NMR measurements of hydrogen/deuterium exchange (HDX) rates reveal a slow unfolding process measured in days. Long pUGs have biphasic unfolding kinetics that include an additional faster phase measured in minutes. The amplitude of this faster phase indicates partial but not global unfolding. These data are consistent with a segmental register exchange model involving flanking sequences on either side of a pUG fold. These data improve our understanding of the structure and folding dynamics of long pUGs involved in gene silencing and many other biological processes.

## Materials and Methods RNA Preparation

RNAs were either chemically synthesized by Integrated DNA Technologies or Dharmacon (Horizon) or prepared by *in vitro* transcription with T7 RNA polymerase. Transcribed RNAs used for NMR experiments 5′-triphosphates were removed with calf-intestinal alkaline phosphatase (Invitrogen) as previously described (11) resulting in all NMR samples having consistent 5′ hydroxyls. All RNAs were purified by denaturant PAGE (8-15% polyacrylamide in 7 M urea and TBE buffer) followed by anion exchange chromatography in Hi-trap Q column (Cytiva). Following purification, RNAs were concentrated and exchanged into pure water using Amicon Ultra 3 kDa or 10 kDa spin filters. Buffer components were added, and the RNA was folded via heating and slow cooling in 1 L of 90 °C water that was allowed to cool to room temperature over the course of 5-6 hrs.

### Oligonucleotide sequences

(GU)12 : 5′-GUGUGUGUGUGUGUGUGUGUGUGU-3′ (GU)13 : 5′-GUGUGUGUGUGUGUGUGUGUGUGUGU-3′

(GU)15 : 5′-GUGUGUGUGUGUGUGUGUGUGUGUGUGUGU-3′

(GU)18 : 5′-GUGUGUGUGUGUGUGUGUGUGUGUGUGUGUGUGUGU-3′

(GU)24 : 5′-GUGUGUGUGUGUGUGUGUGUGUGUGUGUGUGUGUGUGUGUGUGUGUGU-3′

HO-1 pUG: 5′-GUGUGUGUGUGUGUGUGUGUGUGUAUGUGUGUGUGUGUGUGUGUG UGUGU-3′

(GU)29 : 5′-GUGUGUGUGUGUGUGUGUGUGUGUGUGUGUGUGUGUGUGUGUGUGUGUGUGUGUGUGU-3′

U2AA : 5′-GUAAGUGUGUGUGUGUGUGUGUGUGU-3′ U6AA : 5′-GUGUGUAAGUGUGUGUGUGUGUGUGU-3′

### Circular dichroism

Circular dichroism (CD) was recorded using an AVIV model 420 CD spectrometer using quartz cells with a 1 mm path length. CD samples contained roughly 20 μM RNA in 20 mM potassium phosphate buffer, pH 7.0, 100 mM KCl. CD spectra were recorded at 25 °C using a scan step size of 1 nm and 5 s averaging time, measurements were taken from 210 to 340 nm. Raw CD signal was converted to molar CD absorption, Δε.

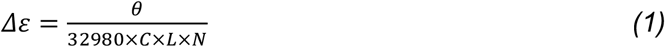

where θ is the raw CD signal (millidegrees), C is RNA concentration (M), L is cuvette path length (cm), and N is number of nucleotides. Thermal denaturation studies utilized a temperature ramp from 20 °C to 81.5 °C in 1.5 °C intervals with a 5 min equilibration time at each temperature.

Ellipticity was measured at four different wavelengths (243, 264 or 304, 284 and 340 nm) with an averaging time of 5 s at each temperature. Melting temperature, Tm, was determined from the thermal unfolding profile at 243, 264 or 304 and 284 nm by fitting to the Boltzmann sigmoidal equation in Origin (Origin 2020, Origin Lab Corporation).

### pUG folding kinetics

Folding was initiated by manually mixing the RNA with 20 mM BIS-TRIS, pH 7.0 buffer, and 150 mM KCl with or without 2 mM MgCl2 to a final concentration of 15-20 μM. Folding kinetics were monitored using CD spectroscopy at four wavelengths corresponding to the characteristic peaks of the pUG fold (243 nm, 264 nm, 284 nm and 304 nm) and one control (340 nm). Ellipticity was measured for 3-5.5 hrs with 100-180 data points with 3 s averaging time. Final CD spectra were collected on all samples to ensure proper folding. Raw CD signal was converted to molar CD absorption (Δε) via equation 1. Data was measured in duplicate to ensure reproducibility. Forward folding half-lives were determined by fitting the Δε vs time curves to a single exponential model using least squares regression in GraphPad Prism. Control experiments containing no salt showed no evidence of pUG fold formation.

### RNase mapping

The (GU)29 RNA was prepared in 50 mM Tris, pH 7.0 with 150 mM KCl or 20 mM LiCl with 20 μM RNA. The samples were folded as described above. RNase treatments were 0.1 U/μL or 0.01 U/μL RNase T1 (Thermo Scientific) for 4 min for the KCl sample and 0.001 U/μL RNase T1 for 1 min for the LiCl sample, both at room temperature (∼20 °C). Treatments were stopped by the addition of an equal volume of 100% formamide. RNase resistant populations were visualized via SYBR Gold nucleic acid gel stain (Invitrogen) stained denaturant PAGE (10% polyacrylamide in 7 M urea and TBE buffer) and fluorescence imaged on an Amersham Typhoon RGB/5 laser-scanner platform. RNase resistant populations were quantified using ImageJ software.

### Native gel analysis

RNA samples were prepared in 20 mM BIS-TRIS, pH 7.0 with 150 mM KCl or LiCl and contained 20 μM RNA. RNA was loaded with 20% sucrose onto a 1.5 mm thick 10% polyacrylamide native electrophoresis gel containing 5 mM KCl and 1X TBE. The gel was run at 5 W for 2.75 hrs at 4 °C with 5 mM KCl in the running buffer. The RNA bands were visualized via staining with toluidine.

### NMR spectroscopy

NMR data was collected at the National Magnetic Resonance Facility at Madison on Bruker Avance III HD 600 MHz or 750 MHz and Varian VNMRS DD 800 MHz spectrometers all equipped with cryogenic probes. The temperature of all samples was regulated at 20 °C. NMR samples were prepared in 20 mM potassium phosphate buffer, pH 7.0 and 100 mM KCl. After folding 10% D2O and 20 μM 4,4-dimethyl-4-silapentane-1-sulfanate (DSS) were added. The same set of experiments were recorded for (GU)12, (GU)13, (GU)15 and (GU)18. Two 1D ^1^H spectra were recorded the first being a full spectrum with water suppression using excitation sculpting with gradients the second using a gradient echo with a selective proton pulse centered on the imino protons, referred to henceforth as a 1D SOFAST. The 2D ^1^H,^1^H NOESY spectra collected in H2O used a 2D version of a SOFAST-NOESY ^15^N-HMQC experiment (22) optimized for imino protons with a 200 ms mixing time. Samples were lyophilized and transferred to 100% D2O and 2D ^1^H,^1^H NOESYs with 200 ms mixing times were recorded to observe the NOEs between non-exchangeable protons. 1D ^31^P spectra were recorder using power-gated decoupling. 1D spectra were processed using MNova software and chemical shifts were referenced to the DSS chemical shift standard set to 0 ppm. The 2D spectra were processed using NMRPipe (23) and analyzed using NMRFAM-Sparky (24).

### Hydrogen/deuterium exchange NMR

All hydrogen/deuterium exchange (HDX) experiments were conducted following the same procedure. RNA samples with concentrations between 100 and 250 μM were prepared in NMR buffer in 90% H2O:10% D2O. Pre-exchange 1D ^1^H and 1D SOFAST spectra centered on the imino region were collected. The sample was removed from the NMR tube and lyophilized to remove solvent. HDX experiment is initiated by the rapid addition of 300 μL of 99.99% D2O at which point the time was noted and data collection began immediately with the first spectra collected within 15-20 min of initiation. Imino proton signal was observed using 1D SOFAST spectra which were continuously recorded for approximately 3 days with additional spectra recorded over the following days/weeks. All spectra were recorded at 20 °C and used the same acquisition parameters. A sample temperature of 20 °C was maintained throughout the course of the experiment. Spectra were processed in MNova and peak integrations were recorded. Integral values were normalized relative to the first data point collected. Data was fit using least squares regression in GraphPad Prism to Equation 2 to determine the observed exchange rate, k_ex_ (25).

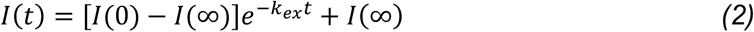

Where I(t) is peak intensity measured as normalized absolute integrals at exchange time t. I(∞) is the intensity of a fully exchanged sample which can be non-zero due to residual H2O in the sample, this value was constrained to 0 as no imino proton signal was observed after months in D2O indicating the residual H2O to be negligible. For (GU)12 the well resolved peaks could be fit separately or grouped by quartet with the fit of the quartet being approximately the same as the average fit of the individual peaks (Table 2). For the longer pUGs, where broadened imino signal made integrating single peaks impossible, the entire imino region was integrated together and fit simultaneously and compared to that of (GU)12 fit in the same manor (Table 3). To verify the validity of this method for analyzing the data, integrations of imino region were also split in half and fit separately to ensure the global fit agreed with the regional fit (Supplemental Figure 1, Supplemental Table 1).

**Table 1:** pUG RNA fraction folded 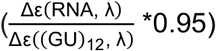 by length

| RNA | $\lambda=245$ nm | $\lambda=281$ nm | $\lambda=304$ nm | $^{24}/_{\text{total nt}}$ |
| --- | --- | --- | --- | --- |
| (GU)<sub>12</sub> | 0.95 ± 0.04 | 0.95 ± 0.02 | 0.95 ± 0.03 | 1 |
| (GU)<sub>13</sub> | 0.92 ± 0.04 | 0.91 ± 0.01 | 0.95 ± 0.01 | 0.92 |
| (GU)<sub>15</sub> | 0.75 ± 0.01 | 0.75 ± 0.01 | 0.68 ± 0.02 | 0.80 |
| (GU)<sub>18</sub> | 0.66 ± 0.02 | 0.69 ± 0.06 | 0.61 ± 0.04 | 0.67 |
| (GU)<sub>29</sub> | 0.74 ± 0.03 | 0.71 ± 0.01 | 0.71 ± 0.004 | 0.41 |

**Table 2:** (GU)12 HDX results

|  | I(0) | k<sub>ex</sub> (hr<sup>-1</sup>) | Half Life (hr) | R squared |
| --- | --- | --- | --- | --- |
| G9/15 | 0.99 ± 0.01 | 0.0054 ± 0.0003 | 127.4 ± 5.8 | 0.976 |
| G3/21 | 0.82 ± 0.02 | 0.0064 ± 0.0007 | 108.8 ± 11.5 | 0.873 |
| G17 | 0.94 ± 0.01 | 0.0114 ± 0.0005 | 60.7 ± 2.5 | 0.978 |
| G5/11 | 0.97 ± 0.01 | 0.0125 ± 0.0004 | 55.7 ± 1.6 | 0.991 |
| G23 | 0.96 ± 0.02 | 0.0179 ± 0.0007 | 38.8 ± 1.4 | 0.981 |

**Table 3:** pUG HDX results length comparison

|  | I(0) | k<sub>ex,fast</sub> (hr<sup>-1</sup>) | k<sub>ex,slow</sub> (hr<sup>-1</sup>) | t<sub>1/2,slow</sub> (hr) | t<sub>1/2,fast</sub> (hr) | Ratio Fast (%) | R-squared |
| --- | --- | --- | --- | --- | --- | --- | --- |
| (GU)<sub>12</sub> | 0.92 ± 0.01 | - | 0.009 ± 0.0004 | 74.8 ± 3.5 | - | - | 0.976 |
| (GU)<sub>13</sub> | 1.14 ± 0.03 | 1.34 ± 0.08 | 0.008 ± 0.0002 | 92.1 ± 1.8 | 0.52 ± 0.03 | 50.2 ± 1.1 | 0.998 |
| (GU)<sub>15</sub> | 1.08 ± 0.04 | 1.29 ± 0.13 | 0.005 ± 0.0004 | 148.6 ± 11.2 | 0.54 ± 0.05 | 52.4 ± 1.6 | 0.986 |
| (GU)<sub>18</sub> | 1.05 ± 0.04 | 1.01 ± 0.12 | 0.004 ± 0.0003 | 171.8 ± 12.2 | 0.69 ± 0.08 | 50.8 ± 2.0 | 0.978 |

### Molecular Dynamics (MD) Simulation

MD simulation was performed with GROMACS v.2023 (26) within the NMRbox resource (27) using the AMBER forcefield with χOL3 modifications (28). The (GU)24 structure was modeled by joining the 5′ and 3′ ends of two copies of the (GU)12 model shown in Figure 1. The (GU)12 model was generated from the solution structure of the pUG fold (GU)12 (PDB ID: 8TNS) and the crystal structure (PDB ID: 7MKT). The quartet rmsd between these models is 0.6 Å, and the combination of them allows for visualization of the ions and all nucleotides including the 3′ terminal U24 which is not observed in the crystal structure due to disorder. The (GU)24 RNA model was solvated in a cubic box of TIP3P water molecules, with a minimum distance of 1 nm from the RNA to the box edges. K^+^ ions were added to a final concentration of 140 mM to neutralize the system. Energy minimization was performed over 50000 steps using the steepest descent algorithm. Next, 100 ps of NVT (number of particles, volume, and temperature) equilibration was applied over 50000 steps using 2 fs timesteps and a modified Berendsen thermostat that gradually increased temperature from 0 to 300 K. This was followed by 100 ps of NPT (number of particles, pressure, and temperature) equilibration using 50000 steps and 2 fs timesteps with a modified Berendsen barostat with pressure maintained at 1 bar. Long-range electrostatic interactions were calculated using the Particle Mesh Ewald method. A production run of 3000 ns was performed with 2 fs timesteps, and trajectory coordinates were saved every 10 ps. Trajectory analysis was carried out using the GROMACS, and visualized with PyMOL v3.1.1 Schrödinger, LLC.

## Results

### Long pUG sequences contain multiple pUG folds

Humans have >20,000 pUGs with 12 or more repeats (24 nucleotides), and among these the average length is 17 repeats and the maximum length is 61 repeats (Supplemental Figure 2). There are >1500 pUGs sequences with 24 or more repeats in human genes (Supplemental Figure 2). In *C. elegans*, long pUG tails >18.5 repeats are more active for gene silencing (11,20). We therefore investigated the structure and folding of long pUG RNAs. The pUG fold is a surprisingly complex structure despite its simple sequence, with 4 different layers of quartets, each in a different conformation with respect to sugar pucker, glycosidic bond angle and backbone orientation (Figure 1). These quartets are: G1-7-13-19, G3-9-15-21, U4-U10-U16-U22 and G5-11-17-23 and hereafter are referred to as the G1, G3, U4 and G5 quartets respectively (Figure 1). The circular dichroism (CD) spectrum of the pUG fold is rich in structural information and reports the conformation of stacked nucleobases and the handedness of the backbone (11). The pUG fold CD spectrum has 4 peaks (Figure 2), which can be attributed to the different types of stacking and backbone orientations in the structure. The negative 245 nm peak and positive 260 nm peaks are representative of right-handed parallel *anti-anti* stacking interactions observed in many G4s (29). The right-handed *anti-anti* interactions in the pUG fold occur at the U4-G5 quartets (Figure 1D). The 260 nm peak is not unique to G4s and is also a feature of right-handed *anti-anti* stacking in A-form RNA (30). The negative peak at 304 nm is unusual and reports on the left-handed Z-form *syn-anti* stacking (30,31) observed at the G3-G5 quartets. Finally, we can attribute the remaining positive peak at 281 nm to the unusual *syn-syn* stacking interactions of the G1-G3 quartets.

**Figure 2.**
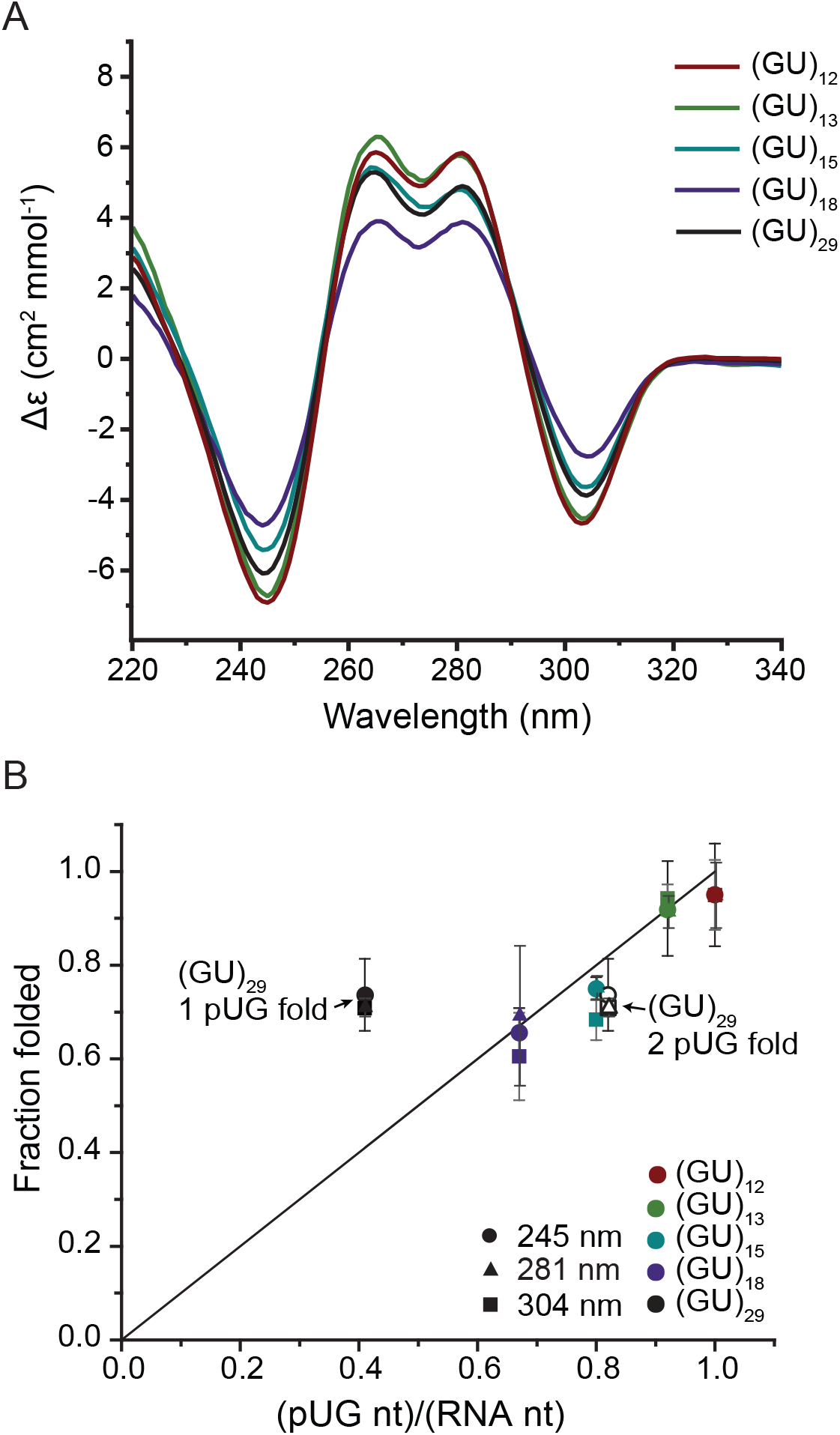
(A) Circular dichroism spectra of (GU)12, (GU)13, (GU)15, (GU)18 and (GU)29. (B) Plot of fraction folded vs nucleotides in a pUG fold per total nucleotides, (pUG nt)/(RNA nt), from the relative magnitudes of the CD signals (Δε(RNA)/Δε((GU)12)*0.95) at 245 nm (circle), 281 nm (triangle) and 304 nm (square). Two points for (GU)29 depict a folding ratio for 1 pUG fold (filled markers) or 2 pUG folds (open markers).

We used CD to investigate the folded state of pUG RNAs of various lengths by heating them to 90°C in 130 mM K^+^ buffer and allowing them to slowly cool over the course of 5 hours. The CD spectra reveal they all adopt pUG folds (Figure 2A). The molar CD absorption (Δε) also reports the fraction of the RNA chain that adopts the pUG fold. We analyzed the CD peaks at 245, 281 and 304 nm for pUG RNAs of different lengths, which reveals a near-perfect linear trend (R^2^=0.99) for fraction folded vs. length (Figure 2). The minimal pUG fold (GU)12 (Figure 1) displays the greatest molar CD absorption, as all 24 nucleotides are engaged in the pUG fold except for the unstructured 3′ terminal U24 (11) (Figure 2A). As the chain lengthens, the molar CD absorption decreases as the fraction of folded nucleotides decreases (Figure 2B). This linear trend indicates an increasing amount of single stranded (ssRNA) content as the chain grows from 12-18 repeats. ^31^P NMR spectra of these RNAs also show the same increase in ssRNA content, observed as a peak of increasing intensity at -0.75 ppm (Supplemental Figure 3). From integration of the ^31^P NMR data, we can normalize the fraction folded for (GU)12 to 95%. Using the normalized molar CD absorption, we can determine the fraction folded of all RNAs; for example, (GU)18 is 65% folded and 35% unstructured (Table 1). The fraction folded agrees well with one pUG fold per RNA for all pUGs ranging from 12-18 repeats (Table 1). The 58-nucleotide pUG RNA (GU)29 deviates markedly from this linear relationship. (GU)29 is long enough to adopt 2 pUG folds and indeed, accounting for 2 pUG folds fits the observed linear relationship of fraction folded (Figure 2B). Two pUG folds in (GU)29 would engage 24 repeats (48 out of 58 nucleotides), corresponding to 83% folded, while a single pUG fold would correspond to only 41% folded. From the measured molar CD absorption, (GU)29 is ∼73% folded, corresponding to an average of ∼1.76 pUG folds per 58 nucleotides. Therefore, the majority of the (GU)29 RNAs have 2 pUG folds, but some have only one.

**Figure 3.**
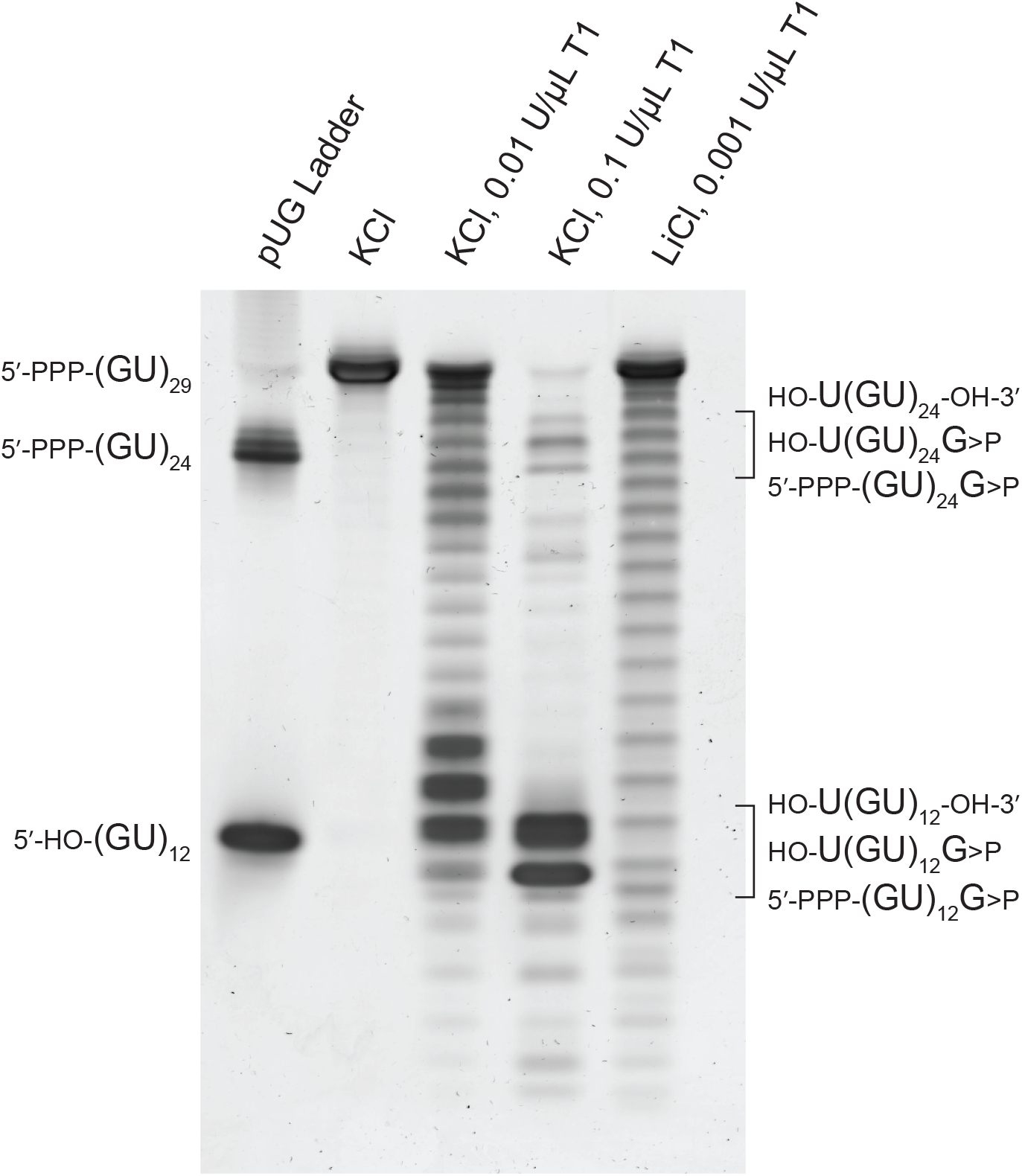
Nuclease mapping of (GU)29 using RNase T1. The first lane contains markers for (GU)12 and (GU)24 RNAs, while the second lane contains untreated (GU)29 in K^+^ buffer. RNase treatments are indicated. The three possible RNase resistant fragments are indicated and correspond to 5′ hydroxyl (HO-), 5′ triphosphate (PPP), 3′ hydroxyl and 3′ cyclic phosphate (>P) termini containing cleavage products.

To further test the idea that multiple pUG folds can form in long pUG RNAs, we subjected (GU)29 to extensive degradation with RNase T1, which cuts after single stranded guanosines, and observed the degradation patterns after staining with SYBR-gold nucleic acid stain (Figure 3). After complete digestion of the intact RNA, we observe RNase-resistant fragments that correspond to single and double pUG folds with 12 and 24 repeats, respectively (Figure 3). In this experiment, RNAs with one pUG fold or two pUG folds separated by unstructured nucleotides are digested into single pUG folds, while compact, adjacent double pUG folds with no intervening nucleotides remain inaccessible to RNase T1. The electrophoretic mobilities of the cleavage products differ slightly depending upon their termini and the position of the pUG fold in the sequence (for example, a pUG fold at the 5′ end has a 5′ guanosine triphosphate, while pUG folds derived from the middle of the RNA have 5′ hydroxy uridines). The adjacent double pUG fold fragments account for ∼18% of the observed cleavage products. This correlates well with the expected distribution for a stochastic folding model, which predicts 22% of all possible folds should result in adjacent double pUG folds (Supplemental Figure 4). The stochastic positioning *in vitro* is consistent with previous chemical probing results of single pUG folds in short RNAs (32).

**Figure 4.**
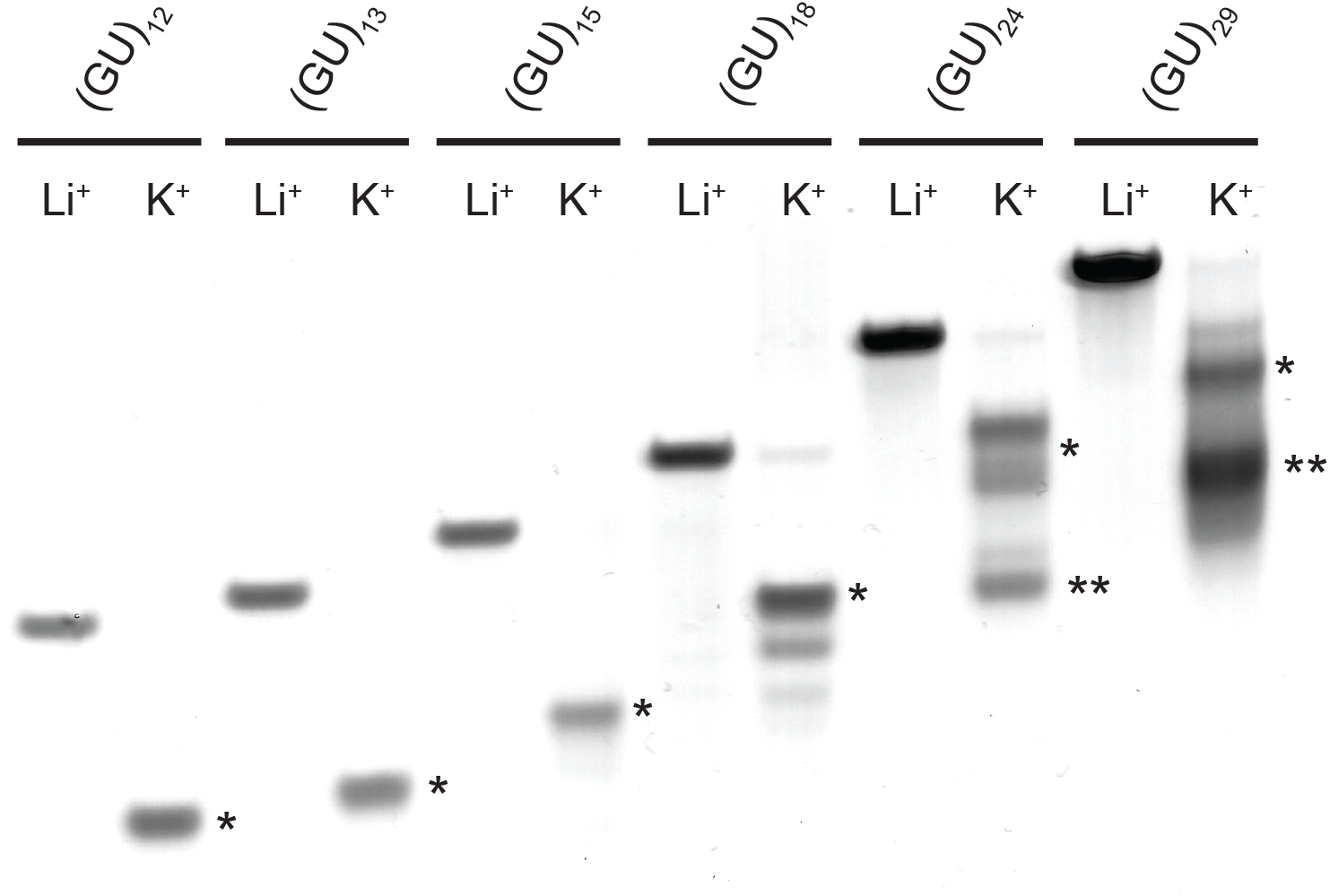
Native PAGE analysis of pUG RNAs of various lengths in K^+^ or Li^+^. Single pUG folds and double pUG folds are marked with one or two asterisks, respectively.

We analyzed the conformations of different lengths of pUG RNAs by native PAGE. The pUG RNAs with 12-18 repeats all adopt single pUG folds (Figure 4). For longer pUG RNAs, the single pUG folds have heterogenous electrophoretic mobilities, which is likely reflective of their different positions within the RNA. The longer pUGs (GU)24 and (GU)29 both adopt single and double pUG folds (Figure 4). (GU)29 is predominately in a double pUG fold conformation, which agrees well with CD data. The faster electrophoretic mobility of the double pUG fold conformation during native PAGE (Figure 4) is consistent with the compact, adjacent double pUG fold conformation identified by nuclease digestion experiments (Figure 3). Our previously determined x-ray and NMR structures of the single pUG fold show the 5′ and 3′ ends are closely positioned, but the 3′ terminal uridine is flexible (11,13). Nevertheless, it is not immediately apparent how adjacent double pUG folds form and avoid steric clash. We therefore used our previously determined structures (11,13) to model the adjacent double pUG fold conformation, which we analyzed by molecular dynamics simulation for 3 μs (Figure 5). While the initial model was in a side-by-side configuration, a stable stacked conformation formed after 485 ns and remained stable for the rest of the 3 μs simulation. Collectively, these data indicate that compact double pUG folds can form within a long pUG RNA.

**Figure 5.**
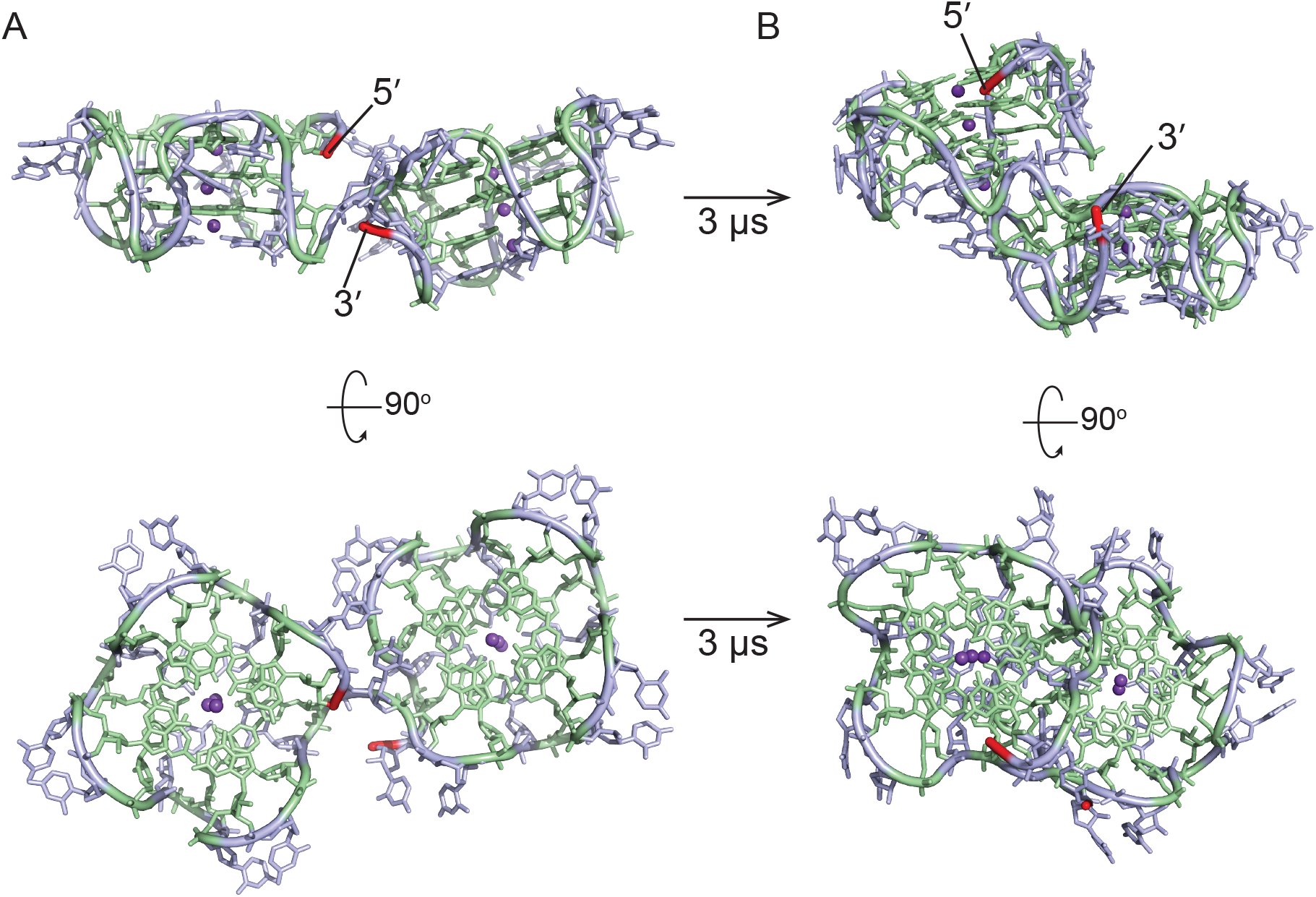
Double pUG fold model of (GU)24. Gs are green, Us are blue, and potassium ions are purple. 5′ and 3′ ends are highlighted in red. (A) depicts the initial conformation following energy minimization and (B) depicts the stacked conformation observed from 486 ns to 3 μs of MD simulation.

### The kinetics of pUG folding

The folding kinetics of pUG RNAs ranging from 12-18 repeats were measured at 25 °C using CD following addition of folding buffer with 150 mM K^+^ or 150 mM K^+^ and 2 mM Mg^2+^. All RNA samples contained 20 μM RNA. The RNAs fold with single exponential kinetics, indicating a monophasic transition from an unfolded to a folded state (Supplemental Figure 5). In 150 mM K^+^ buffer, the folding half-life of (GU)12 is 17 min, (GU)15 is 13 min, and the longer pUG RNA (GU)18 has the slowest folding rate with a half-life of 30 min. The slow folding of the pUG fold is likely a result of its complex structure which includes 8 backbone inversions and 8 *syn* nucleotide conformations (Figure 1D). The kinetics of double pUG folding were not measured but are likely even slower. One day after a 5 hour folding protocol of heating and slow cooling, 78% of (GU)29 contained 2 pUG folds (Figure 2B). After storage for an additional 1 month at 4 °C, 98% of (GU)29 contained 2 pUG folds (Supplemental Figure 6). Thus, long pUG RNAs slowly resolve into double pUG folds over the course of hours to days *in vitro*.

**Figure 6.**
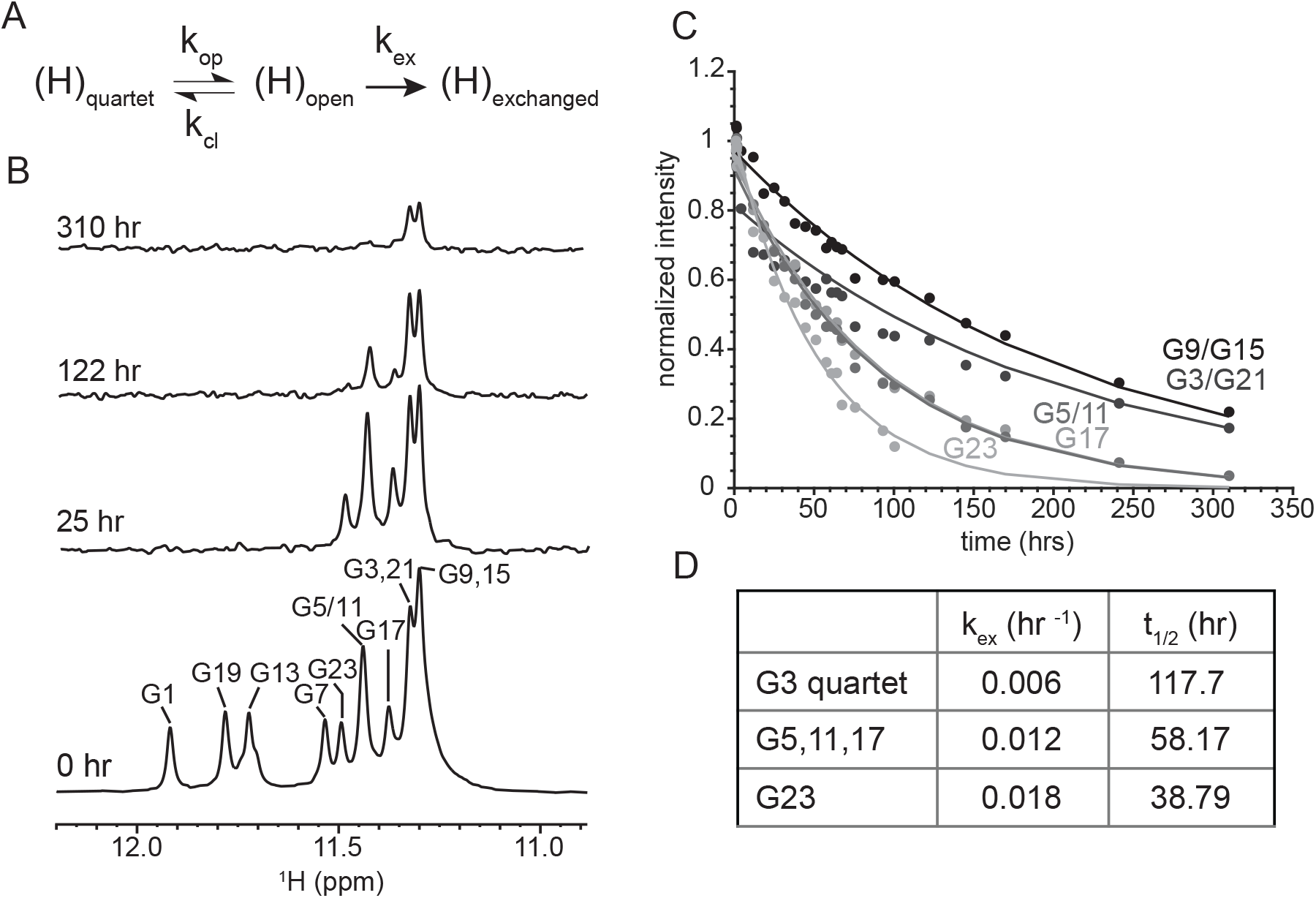
Hydrogen/deuterium exchange (HDX) NMR of (GU)12. (A) HDX mechanism where H-bonded protons in quartets, (H)quartet, must open, (H)open, to allow the proton to be exchanged, (H)exchanged, with deuterium. (B) 1D ^1^H NMR showing loss of imino proton signal at given times following transfer into D2O. (C) Exponential decay of the normalized peak integrals (I(t)/I(0)), over time. (D) Summary of average HDX rates (kex) and quartet half-lives for the G3 and G5 quartets, G23 was measured separately.

Although Mg^2+^ is not required for the pUG fold, we observe that addition of 2 mM Mg^2+^ increases the rate of folding of the minimal pUG fold (GU)12, decreasing the half-life from 17 to 13 min (Supplemental Figure 5). On the other hand, the folding of longer pUGs is inhibited by the addition of Mg^2+^, as (GU)15 folds with a half-life of 36 min in K^+^ and Mg^2+^ buffer and the folding of (GU)18 was incomplete after 5 hours in K^+^ and Mg^2+^ buffer. The CD spectra of (GU)18 in K^+^ and Mg^2+^ buffer shows a positive peak at 260 nm, but the pUG fold peaks at 243, 284 and 304 nm are of much lower intensity (Supplemental Figure 7). These data are consistent with an A-form helical conformation involving GU wobble pairs competing with pUG folding in longer RNAs. Intermolecular association of long pUGs may be facilitated by the relatively high concentrations (20 μm) needed for the CD measurements, as intermolecular A-form helices are stabilized by Mg^2+^ and tandem GU wobble pairs create K^+^ binding sites (33). Nevertheless, (GU)18 in K^+^ and Mg^2+^ buffer still adopts the pUG fold following a heat and slow cool cycle (Supplemental Figure 7).

**Figure 7.**
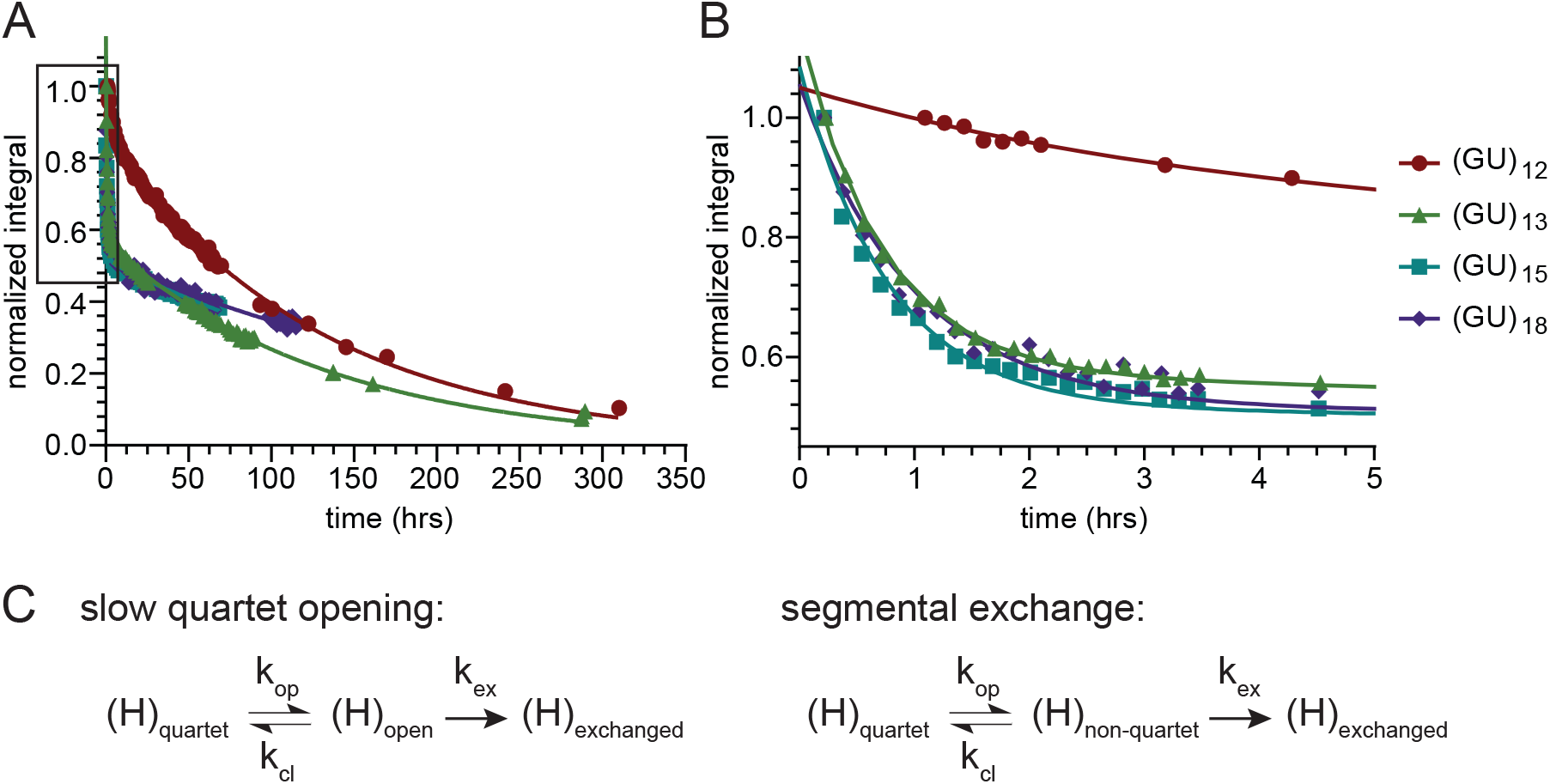
Biphasic HDX of (GU)13, (GU)15 and (GU)18 compared to (GU)12. (A) Plot of loss of imino proton signal measured as normalized integrals (I(t)/I(0)) of the entire imino region over time fit to an exponential decay for (GU)12 and biexponential decays for (GU)13, (GU)15 and (GU)18. (B) Zoomed in plot of boxed region from (A) highlighting the fast phase of HDX biexponential decay for (GU)13, (GU)15 and (GU)18. (C) Mechanism of slow and fast phase corresponding to quartet opening and segmental exchange dynamics respectively.

We allowed pUG RNAs to completely fold in K^+^ buffer and measured the kinetics of quartet opening rates and pUG unfolding using hydrogen/deuterium exchange (HDX) NMR. Upon deuterium solvent exposure, the hydrogen bonded imino proton signals disappear over time (Figure 6)(25,34-36). The solvent exposed G1 and U4 quartets exchange rapidly and were not measured, but the imino signals from the buried G3 and G5 quartets are observable for days in D2O (Figure 6B). By fitting the exponential decay of the imino proton signals over time (Figure 6C), we determined the exchange rates (kex) for the G3 and G5 quartets (Figure 6D, Table 2), which have half-lives of 5 and 2.5 days, respectively. The slow exchange rate of central quartets inform on the overall G4 unfolding rate (37). The G23 guanosine is adjacent to the 3′ end and has a slightly faster exchange rate than the other Gs, which can be attributed to increased dynamics at the more flexible 3′ end (13). The imino proton signals of (GU)13, (GU)15 and (GU)18 were also observable for days after transfer into D2O (Supplemental Figure 8). However, the longer pUGs exhibit biphasic HDX rates (Figure 7) with slow and fast exchange processes (kex,slow and kex,fast) (Table 3). The slow phase corresponds to the slow quartet opening observed in (GU)12 with half-lives in the range of 3.8-7 days (Figure 7C). The faster exchange process corresponds to half-lives of approximately 30 min. The amplitude of the fast phase is ∼50%, which indicates partial but not complete unfolding. An increase in line broadening of NMR spectra is also observed as pUG length increases, which is indicative of dynamics on the microsecond to millisecond timescale (Figure 8 and Supplemental Figure 9). The increase in line broadening is not due to the size of the RNAs, as a larger 50 nucleotide RNA sequence (GU)12AU(GU)12 derived from the 5′ untranslated region of the human gene heme oxygenase 1 (HO-1) (38) has narrower linewidths than the 36 nucleotide (GU)18 (Figure 8). Despite the line broadening, the pUG fold can be identified in 1D (Figure 8) and 2D ^1^H,^1^H NOESY spectra for all RNAs (Supplemental Figure 9). Finally, longer pUG folds also have slightly lower thermal stabilities. The minimal (GU)12 pUG fold has a melting temperature of 51.5 ºC (11), while (GU)13, (GU)15 and (GU)18 have melting temperatures of 48.3 ºC, 47.1 ºC and 47.9 ºC, respectively (Supplemental Figure 10). From these data we conclude that long pUG RNAs form dynamic pUG folds that are kinetically stable and can persist for days after folding.

**Figure 8.**
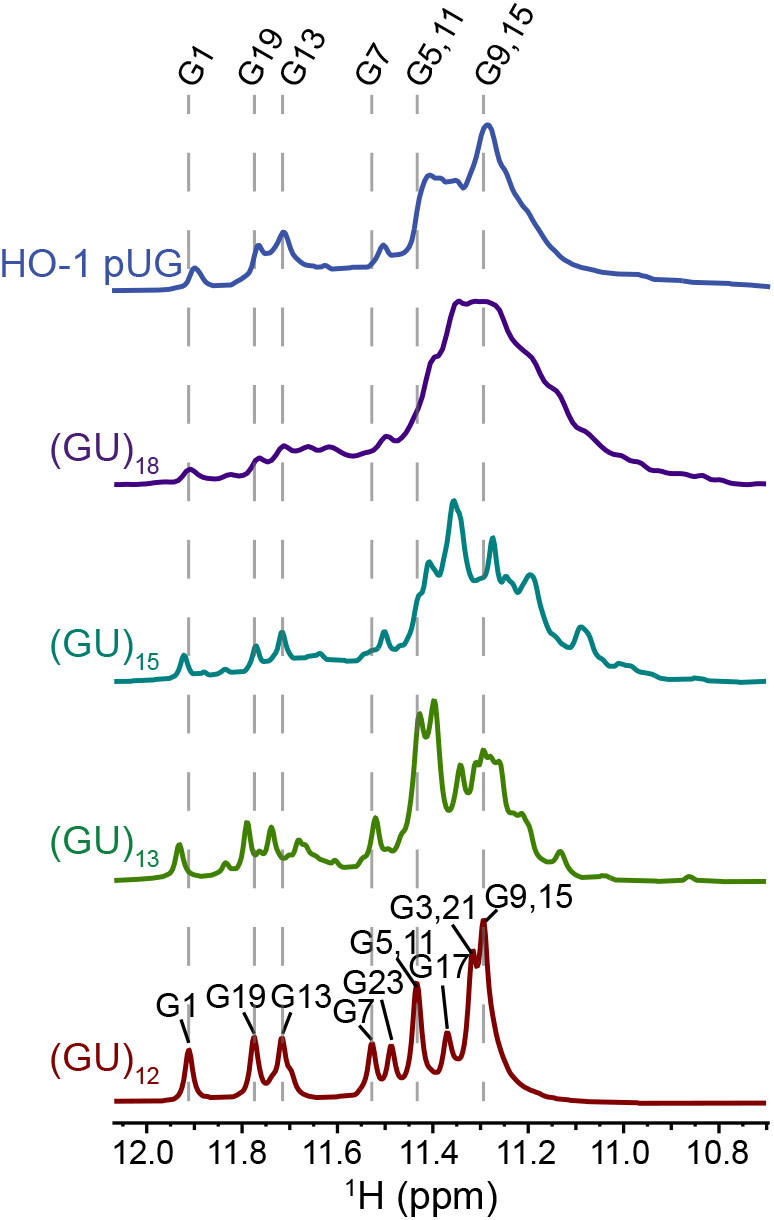
1D ^1^H NMR of (GU)12, (GU)13, (GU)15, (GU)18 and the 50 nt sequence (GU)12AU(GU)12 from the HO-1 gene. Dotted lines mark chemical shifts of the (GU)12 pUG fold.

**Figure 9.**
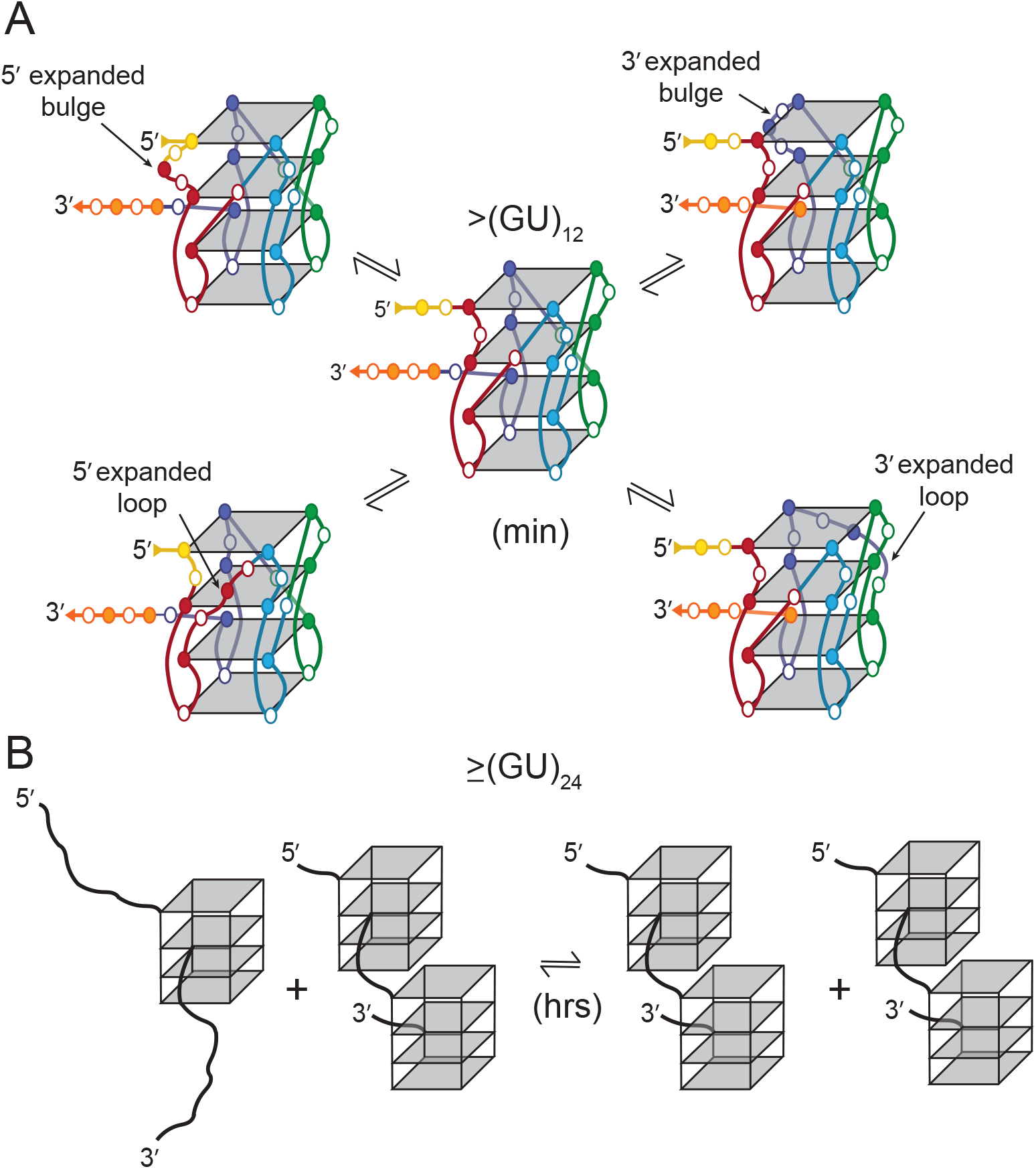
Schematic models of pUG fold dynamics. (A) Segmental register exchange of an internal pUG fold in (GU)15 involving flanking 5′ and 3′ segments. Gs shown in filled circles and Us shown as open circles. Single stranded nucleotides (yellow and orange) exchange into the structure through the formation of expanded bulges or propeller loops. (B) Global structural rearrangement of (GU)29 resulting in an increase in double pUG folds.

## Discussion

Here we show that long pUG RNAs with 24 or more repeats can form compact double pUG folds. This is the first experimental evidence for adjacent double pUG folds. Such long pUG repeats are found in thousands of human RNAs. A well-studied example is the lncRNA NEAT1, which has 29.5 pUG repeats. Dysregulation of NEAT1 has been found in almost all cancers and is associated with chemotherapy resistance and poor prognosis (15-18). The human chromatin modifying enzyme DNMT1 binds to, and is inhibited by, pUG folds (39), and also interacts with NEAT1 (40,41). These data suggest pUG folds can form in RNAs with long pUGs and may contribute to disease. Many human pUG repeats are polymorphic with expansions that correlate with higher incidence of disease. For example, a polymorphic pUG is found in the 5′ untranslated region (UTR) of heme oxygenase 1 (HO-1). HO-1 is the enzyme responsible for breaking down heme and must be tightly regulated within a certain cytoprotective range (38). The pUG repeats in HO-1 are bimodal with short and long isoforms centered on 23 and 30 repeats respectively (42). The long repeat isoform (≥ 25 repeats) has been shown to correlate to higher rates of several diseases such as cardiovascular disease, sickle cell anemia, rheumatoid arthritis and emphysema (6,43-45). Repeat length has also been shown to be an independent prognosticator for several cancers including esophageal and pancreatic (46,47). The long repeat isoform is associated with the use of an alternative first exon which contains an alternative transcription start site leading to lower overall expression of active protein (38). HO-1 expression is stimulated by hemin, but hemin also induces expression of 5′ elongated transcripts (38). Our data suggests that the formation of a second pUG fold in the long repeat isoform may contribute to the mis-splicing of HO-1 or interfere with hemin induced upregulation. Intriguingly, we have previously shown that the pUG fold binds hemin, opening a possible avenue in which HO-1 mRNA may be able to sense hemin to upregulate expression (11).

Although the initial formation of pUG folds *in vitro* is stochastic with respect to their position in the RNA, long pUGs slowly resolve into double pUG folds. Folding *in vivo* may be facilitated by co-transcriptional folding and/or pUG RNA binding proteins. The ability to form double pUG folds provides a potential explanation as to why longer pUG tails are more active for gene silencing *in vivo* (11). The 3′ end of the pUG fold is flexible, as shown by NMR (13) and the MD simulations reported here and allows the formation of closely packed multimeric pUG folds, like beads on a string. Consistent with the idea of multimeric pUG interactions, pUG repeats are more efficient at phase separation than other G-rich sequences (48). Stacking interactions between pUG folds likely contribute to condensation and phase separation (48).

The folding and unfolding kinetics reported here are slower than many RNA tertiary structures (49) but are typical of G4s. For example, the *c-myc* DNA G4 has an average lifetime of ∼2 days (35,36) while the pUG fold is ∼7 days. Our HDX data reveal that pUG RNAs with more than 12 repeats display an additional dynamic phase that is much faster than global unfolding. The amplitude of this faster phase is ∼50%, consistent with partial but not global unfolding. A model depicting such a semi-conservative conformational exchange process is illustrated (Figure 9A). This model involves the invasion of flanking sequences to induce local register exchange of the 5′ and 3′ segments, accompanied by the formation of expanded loops or bulges (Figure 9A). Such a dynamic rearrangement explains the 50% amplitude of the fast phase and the fact that this process is not observed for the minimal pUG fold with only 12 repeats. To further test the validity of this model, we created pUG folds with targeted AA insertions within the U2 bulge or the U6 loop to enforce the formation of larger loops, and these RNAs still adopt pUG folds (Supplemental Figure 11). Dynamic formation of larger loops is also consistent with the decrease in thermodynamic stability measured for longer pUGs which is also observed for other G4 structures with loops (50). The dynamic model involving 5′ and 3′ flanking segments implies that structural heterogeneity should increase with pUG length, as longer flanking segments can pair in different registers. Indeed, we observe the presence of additional peaks in the ^1^H NMR spectra of longer pUGs (Figure 8). These additional peaks are consistent with heterogeneity in loops and bulges but may also arise from other conformations. For example, the pUG fold can be formed through association of two short (GU)6 strands that interact *in trans* to form a dimeric pUG fold (48). Thus, pUGs with 18 or more repeats may be able to simultaneously form both intramolecular and intermolecular pUG folds.

Despite dynamic conformational exchange, the pUG fold core is stable for days. The kinetic stability of the pUG fold may provide a protective function in *C. elegans*, where pUG tails are RNA epigenetic modifications that are passed on for generations. The pUG fold must recruit RdRP to these templates, while protecting them from degradation by the 3′-5′ exoribonuclease MUT-7 which is also essential for gene silencing in *C. elegans* (51). A proposed role of MUT-7 is to trim 3′ overhanging nucleotides from the pUG fold (51), which could lead to homogenous pUG folds by preventing the formation of loops.

In this work we have measured the folding and unfolding rates of pUG RNAs of varying lengths. We show that pUG folds have dynamic folding pathways and can form compact, multimeric double pUG folds. The relatively slow unfolding dynamics of pUGs is typical of G4 structures (34). The persistence of the pUG fold for days is likely due to extensive stacking and potassium ion coordination. Kinetic stability may be important *in vivo*, as pUG fold-containing RNAs are packaged in embryos and inherited for generations in *C. elegans* (11,20). The pUG fold may provide a unique, left-handed platform for proteins (39) and small molecule interactions (11). Finally, in humans there are thousands of pUGs, many of which have been associated with disease, suggesting important biological roles that are yet to be discovered.

## Data Availability

The data underlying this article will be shared on reasonable request to the corresponding author.

## Supplementary Data

Supplementary Data are available at BioRxiv.

## Acknowledgements

This study made use of the National Magnetic Resonance Facility at Madison, an NIH Biomedical Technology Research Resource Center NIH R24GM141526. Helium recovery equipment, computers, and infrastructure for data archive were funded by the University of Wisconsin-Madison, NIH R24GM141526. This study made use of NMRbox: National Center for Biomolecular NMR Data Processing and Analysis, a Biomedical Technology Research Resource (BTRR), which is supported by NIH grant P41GM111135 (NIGMS). CD data were obtained at the University of Wisconsin-Madison Biophysics Instrumentation Facility, which was established with support from the University of Wisconsin-Madison and grants BIR-9512577 (NSF) and S10RR013790 (NIH).

## Author Contributions

R.J.P., R.V., M.T. and S.R. designed and performed experiments. R.J.P. analyzed data, interpreted results and wrote the manuscript. S.E.B. interpreted results, wrote the manuscript, and obtained funding.

## References

1. Lander, E.S., Linton, L.M., Birren, B., Nusbaum, C., Zody, M.C., Baldwin, J., Devon, K., Dewar, K., Doyle, M., FitzHugh, W. et al. (2001) Initial sequencing and analysis of the human genome. Nature, 409, 860–921.

2. Toth, G., Gaspari, Z. and Jurka, J. (2000) Microsatellites in different eukaryotic genomes: survey and analysis. Genome Res, 10, 967–981.

3. Groman, J.D., Hefferon, T.W., Casals, T., Bassas, L., Estivill, X., Des Georges, M., Guittard, C., Koudova, M., Fallin, M.D., Nemeth, K. et al. (2004) Variation in a repeat sequence determines whether a common variant of the cystic fibrosis transmembrane conductance regulator gene is pathogenic or benign. Am J Hum Genet, 74, 176–179.

4. Buratti, E., Brindisi, A., Pagani, F. and Baralle, F.E. (2004) Nuclear factor TDP-43 binds to the polymorphic TG repeats in CFTR intron 8 and causes skipping of exon 9: a functional link with disease penetrance. Am J Hum Genet, 74, 1322–1325.

5. Gao, P.S., Heller, N.M., Walker, W., Chen, C.H., Moller, M., Plunkett, B., Roberts, M.H., Schleimer, R.P., Hopkin, J.M. and Huang, S.K. (2004) Variation in dinucleotide (GT) repeat sequence in the first exon of the STAT6 gene is associated with atopic asthma and differentially regulates the promoter activity in vitro. J Med Genet, 41, 535–539.

6. Daenen, K.E., Martens, P. and Bammens, B. (2016) Association of HO-1 (GT)n Promoter Polymorphism and Cardiovascular Disease: A Reanalysis of the Literature. Can J Cardiol, 32, 160–168.

7. Itokawa, M., Yamada, K., Yoshitsugu, K., Toyota, T., Suga, T., Ohba, H., Watanabe, A., Hattori, E., Shimizu, H., Kumakura, T. et al. (2003) A microsatellite repeat in the promoter of the N-methyl-D-aspartate receptor 2A subunit (GRIN2A) gene suppresses transcriptional activity and correlates with chronic outcome in schizophrenia. Pharmacogenetics, 13, 271–278.

8. Tang, J., Chen, X., Xu, X., Wu, R., Zhao, J., Hu, Z. and Xia, K. (2006) Significant linkage and association between a functional (GT)n polymorphism in promoter of the N-methyl-D-aspartate receptor subunit gene (GRIN2A) and schizophrenia. Neurosci Lett, 409, 80–82.

9. Boraska Jelavic, T., Barisic, M., Drmic Hofman, I., Boraska, V., Vrdoljak, E., Peruzovic, M., Hozo, I., Puljiz, Z. and Terzic, J. (2006) Microsatelite GT polymorphism in the toll-like receptor 2 is associated with colorectal cancer. Clin Genet, 70, 156–160.

10. Sawaya, S., Bagshaw, A., Buschiazzo, E., Kumar, P., Chowdhury, S., Black, M.A. and Gemmell, N. (2013) Microsatellite tandem repeats are abundant in human promoters and are associated with regulatory elements. PLoS One, 8, e54710.

11. Roschdi, S., Yan, J., Nomura, Y., Escobar, C.A., Petersen, R.J., Bingman, C.A., Tonelli, M., Vivek, R., Montemayor, E.J., Wickens, M. et al. (2022) An atypical RNA quadruplex marks RNAs as vectors for gene silencing. Nat Struct Mol Biol, 29, 1113–1121.

12. Butcher, S.E. (2024) A left-handed RNA quadruplex directs gene silencing. Trends Biochem Sci, 49, 387–390.

13. Escobar, C.A., Petersen, R.J., Tonelli, M., Fan, L., Henzler-Wildman, K.A. and Butcher, S.E. (2023) Solution Structure of Poly(UG) RNA. J Mol Biol, 435, 168340.

14. Hefferon, T.W., Groman, J.D., Yurk, C.E. and Cutting, G.R. (2004) A variable dinucleotide repeat in the CFTR gene contributes to phenotype diversity by forming RNA secondary structures that alter splicing. Proc Natl Acad Sci U S A, 101, 3504–3509.

15. Gu, J., Zhang, B., An, R., Qian, W., Han, L., Duan, W., Wang, Z. and Ma, Q. (2022) Molecular Interactions of the Long Noncoding RNA NEAT1 in Cancer. Cancers (Basel), 14.

16. Long, F., Li, X., Pan, J., Ye, H., Di, C., Huang, Y., Li, J., Zhou, X., Yi, H., Huang, Q. et al. (2024) The role of lncRNA NEAT1 in human cancer chemoresistance. Cancer Cell Int, 24, 236.

17. Yang, C., Li, Z., Li, Y., Xu, R., Wang, Y., Tian, Y. and Chen, W. (2017) Long non-coding RNA NEAT1 overexpression is associated with poor prognosis in cancer patients: a systematic review and meta-analysis. Oncotarget, 8, 2672–2680.

18. Yong, W., Yu, D., Jun, Z., Yachen, D., Weiwei, W., Midie, X., Xingzhu, J. and Xiaohua, W. (2018) Long noncoding RNA NEAT1, regulated by LIN28B, promotes cell proliferation and migration through sponging miR-506 in high-grade serous ovarian cancer. Cell Death Dis, 9, 861.

19. Knutsen, E., Harris, A.L. and Perander, M. (2022) Expression and functions of long non-coding RNA NEAT1 and isoforms in breast cancer. Br J Cancer, 126, 551–561.

20. Shukla, A., Yan, J., Pagano, D.J., Dodson, A.E., Fei, Y., Gorham, J., Seidman, J.G., Wickens, M. and Kennedy, S. (2020) poly(UG)-tailed RNAs in genome protection and epigenetic inheritance. Nature, 582, 283–288.

21. Preston, M.A., Porter, D.F., Chen, F., Buter, N., Lapointe, C.P., Keles, S., Kimble, J. and Wickens, M. (2019) Unbiased screen of RNA tailing activities reveals a poly(UG) polymerase. Nat Methods, 16, 437–445.

22. Schanda, P. and Brutscher, B. (2005) Very fast two-dimensional NMR spectroscopy for real-time investigation of dynamic events in proteins on the time scale of seconds. J Am Chem Soc, 127, 8014–8015.

23. Delaglio, F., Grzesiek, S., Vuister, G.W., Zhu, G., Pfeifer, J. and Bax, A. (1995) NMRPipe: a multidimensional spectral processing system based on UNIX pipes. J Biomol NMR, 6, 277–293.

24. Lee, W., Tonelli, M. and Markley, J.L. (2015) NMRFAM-SPARKY: enhanced software for biomolecular NMR spectroscopy. Bioinformatics, 31, 1325–1327.

25. Russu, I.M. (2004) Probing site-specific energetics in proteins and nucleic acids by hydrogen exchange and nuclear magnetic resonance spectroscopy. Methods Enzymol, 379, 152–175.

26. Van Der Spoel, D., Lindahl, E., Hess, B., Groenhof, G., Mark, A.E. and Berendsen, H.J. (2005) GROMACS: fast, flexible, and free. J Comput Chem, 26, 1701–1718.

27. Maciejewski, M.W., Schuyler, A.D., Gryk, M.R., Moraru, II, Romero, P.R., Ulrich, E.L., Eghbalnia, H.R., Livny, M., Delaglio, F. and Hoch, J.C. (2017) NMRbox: A Resource for Biomolecular NMR Computation. Biophys J, 112, 1529–1534.

28. Zgarbova, M., Otyepka, M., Sponer, J., Mladek, A., Banas, P., Cheatham, T.E., 3rd and Jurecka, P. (2011) Refinement of the Cornell et al. Nucleic Acids Force Field Based on Reference Quantum Chemical Calculations of Glycosidic Torsion Profiles. J Chem Theory Comput, 7, 2886–2902.

29. Del Villar-Guerra, R., Trent, J.O. and Chaires, J.B. (2018) G-Quadruplex Secondary Structure Obtained from Circular Dichroism Spectroscopy. Angew Chem Int Ed Engl, 57, 7171–7175.

30. Kypr, J., Kejnovska, I., Renciuk, D. and Vorlickova, M. (2009) Circular dichroism and conformational polymorphism of DNA. Nucleic Acids Res, 37, 1713–1725.

31. Hall, K., Cruz, P., Tinoco, I., Jr., Jovin, T.M. and van de Sande, J.H. (1984) ‘Z-RNA’--a left-handed RNA double helix. Nature, 311, 584–586.

32. Oleynikov, M. and Jaffrey, S.R. (2024) RNA tertiary structure and conformational dynamics revealed by BASH MaP. eLife, 13, RP98540.

33. Marcia, M. and Pyle, A.M. (2014) Principles of ion recognition in RNA: insights from the group II intron structures. RNA, 20, 516–527.

34. Hsu, S.T., Varnai, P., Bugaut, A., Reszka, A.P., Neidle, S. and Balasubramanian, S. (2009) A G-rich sequence within the c-kit oncogene promoter forms a parallel G-quadruplex having asymmetric G-tetrad dynamics. J Am Chem Soc, 131, 13399–13409.

35. Phan, A.T., Modi, Y.S. and Patel, D.J. (2004) Propeller-type parallel-stranded G-quadruplexes in the human c-myc promoter. J Am Chem Soc, 126, 8710–8716.

36. Lee, S., Lee, A.R., Ryu, K.S., Lee, J.H. and Park, C.J. (2019) NMR Investigation of the Interaction between the RecQ C-Terminal Domain of Human Bloom Syndrome Protein and G-Quadruplex DNA from the Human c-Myc Promoter. J Mol Biol, 431, 794–806.

37. Li, M.H., Wang, Z.F., Kuo, M.H., Hsu, S.T. and Chang, T.C. (2014) Unfolding kinetics of human telomeric G-quadruplexes studied by NMR spectroscopy. J Phys Chem B, 118, 931–936.

38. Kramer, M., Sponholz, C., Slaba, M., Wissuwa, B., Claus, R.A., Menzel, U., Huse, K., Platzer, M. and Bauer, M. (2013) Alternative 5’ untranslated regions are involved in expression regulation of human heme oxygenase-1. PLoS One, 8, e77224.

39. Jansson-Fritzberg, L.I., Sousa, C.I., Smallegan, M.J., Song, J.J., Gooding, A.R., Kasinath, V., Rinn, J.L. and Cech, T.R. (2023) DNMT1 inhibition by pUG-fold quadruplex RNA. RNA, 29, 346–360.

40. Li, Y. and Cheng, C. (2018) Long noncoding RNA NEAT1 promotes the metastasis of osteosarcoma via interaction with the G9a-DNMT1-Snail complex. Am J Cancer Res, 8, 81–90.

41. Ma, F., Lei, Y.Y., Ding, M.G., Luo, L.H., Xie, Y.C. and Liu, X.L. (2020) LncRNA NEAT1 Interacted With DNMT1 to Regulate Malignant Phenotype of Cancer Cell and Cytotoxic T Cell Infiltration via Epigenetic Inhibition of p53, cGAS, and STING in Lung Cancer. Front Genet, 11, 250.

42. Exner, M., Minar, E., Wagner, O. and Schillinger, M. (2004) The role of heme oxygenase-1 promoter polymorphisms in human disease. Free Radic Biol Med, 37, 1097–1104.

43. Hariharan, P., Chavan, V. and Nadkarni, A. (2020) Significance of heme oxygenase-1(HMOX1) gene on fetal hemoglobin induction in sickle cell anemia patients. Sci Rep, 10, 18506.

44. Rueda, B., Oliver, J., Robledo, G., Lopez-Nevot, M.A., Balsa, A., Pascual-Salcedo, D., Gonzalez-Gay, M.A., Gonzalez-Escribano, M.F. and Martin, J. (2007) HO-1 promoter polymorphism associated with rheumatoid arthritis. Arthritis Rheum, 56, 3953–3958.

45. Yamada, N., Yamaya, M., Okinaga, S., Nakayama, K., Sekizawa, K., Shibahara, S. and Sasaki, H. (2000) Microsatellite polymorphism in the heme oxygenase-1 gene promoter is associated with susceptibility to emphysema. Am J Hum Genet, 66, 187–195.

46. Ghadban, T., Miro, J.T., Trump, F., Tsui, T.Y., Uzunoglu, F.G., Reeh, M., Gebauer, F., Bachmann, K., Wellner, U., Kalinin, V. et al. (2016) Diverse prognostic value of the GTn promoter polymorphism in squamous cell and adeno carcinoma of the oesophagus. Clin Genet, 90, 343–350.

47. Vashist, Y.K., Uzungolu, G., Kutup, A., Gebauer, F., Koenig, A., Deutsch, L., Zehler, O., Busch, P., Kalinin, V., Izbicki, J.R. et al. (2011) Heme oxygenase-1 germ line GTn promoter polymorphism is an independent prognosticator of tumor recurrence and survival in pancreatic cancer. J Surg Oncol, 104, 305–311.

48. Roschdi, S., Montemayor, E.J., Vivek, R., Bingman, C.A. and Butcher, S.E. (2024) Self-assembly and condensation of intermolecular poly(UG) RNA quadruplexes. Nucleic Acids Res, 52, 12582–12591.

49. Woodson, S.A. (2010) Compact intermediates in RNA folding. Annu Rev Biophys, 39, 61–77.

50. Pandey, S., Agarwala, P. and Maiti, S. (2013) Effect of loops and G-quartets on the stability of RNA G-quadruplexes. J Phys Chem B, 117, 6896–6905.

51. Busetto, V., Pshanichnaya, L., Lichtenberger, R., Hann, S., Ketting, R.F. and Falk, S. (2024) MUT-7 exoribonuclease activity and localization are mediated by an ancient domain. Nucleic Acids Res, 52, 9076–9091.

