## SupplementalData for "The structure, folding kinetics and dynamics of long poly(UG) RNA"

### Supplementary Figures

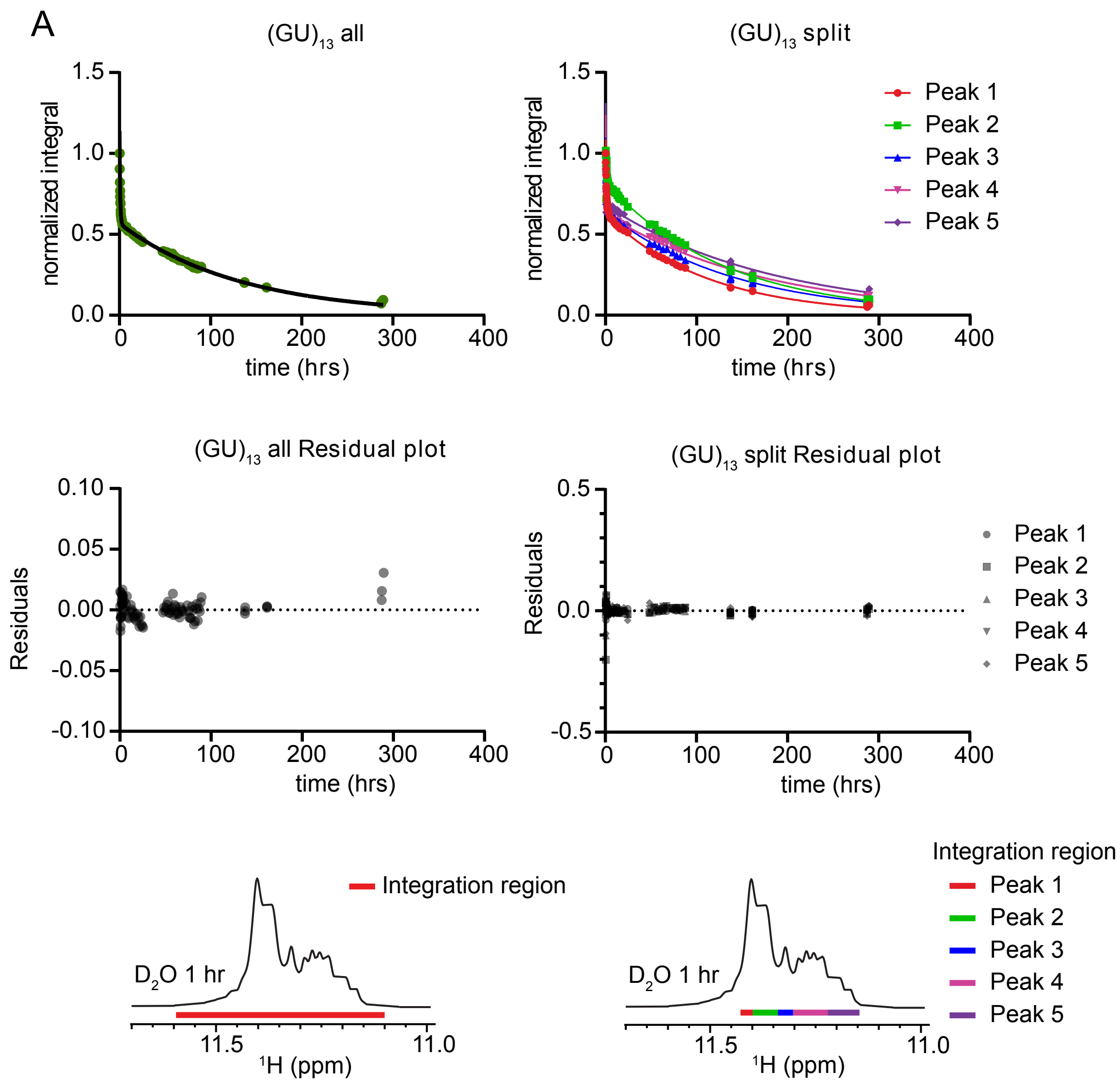


**Supplemental Figure 1A.** Global and regional fits of (A) $(GU)_{13}$, (B) $(GU)_{15}$ and (C) $(GU)_{18}$ HDX. From top to bottom: normalized integral (I(t)/I(0)) vs time plot fit to biexponential decay, residual plots, $^{1}H$ NMR spectrum with integral regions shown. Left column shows entire imino region integrated together to account for broadening, right column shows split integral regions to ensure consistency of fit. Results of fits shown in Supplemental Table 1.

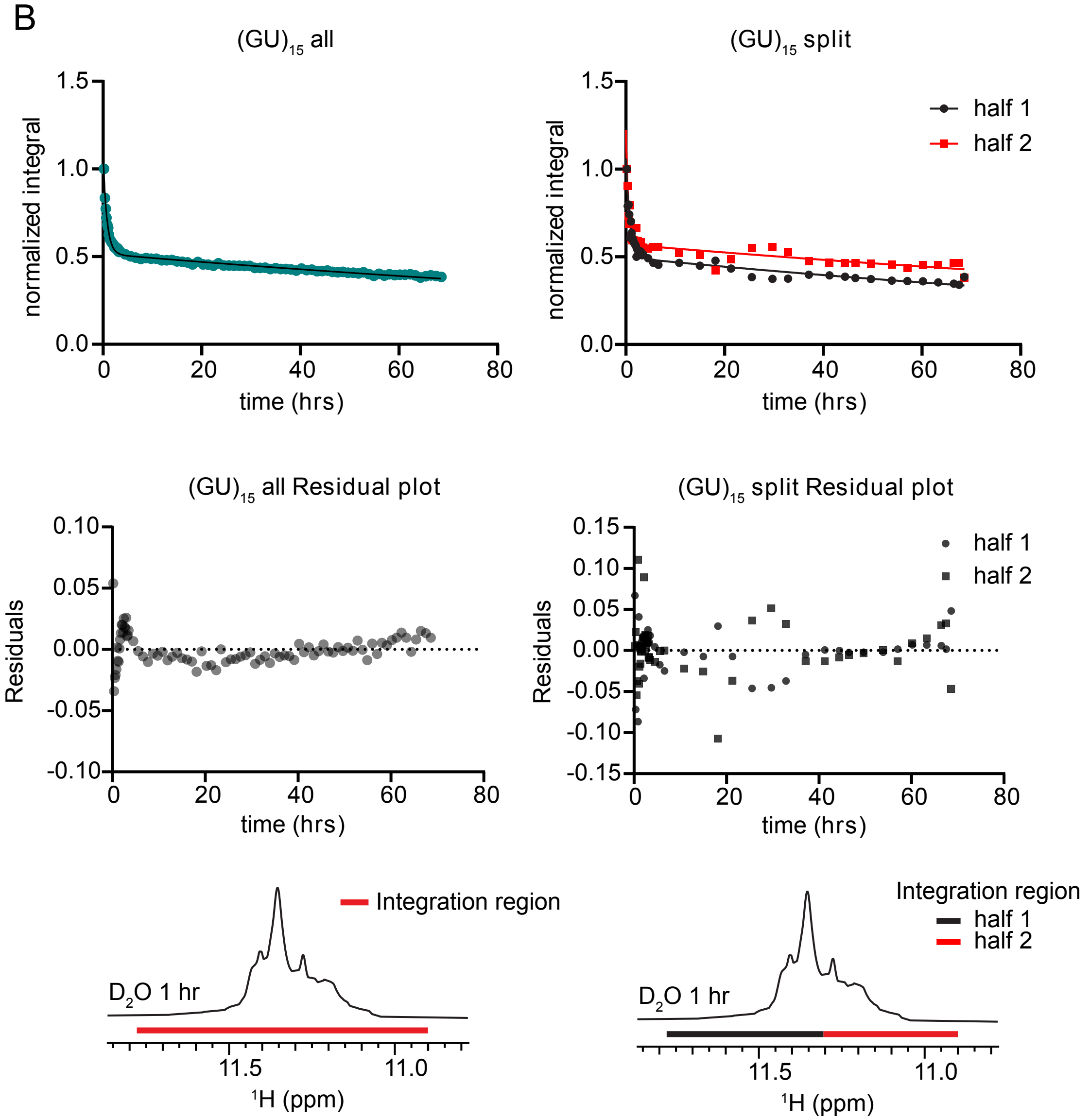


**Supplemental Figure 1B.** Global and regional fits of (A) $(GU)_{13}$, (B) $(GU)_{15}$ and (C) $(GU)_{18}$ HDX. From top to bottom: normalized integral (I(t)/I(0)) vs time plot fit to biexponential decay, residual plots, $^{1}H$ NMR spectrum with integral regions shown. Left column shows entire imino region integrated together to account for broadening, right column shows split integral regions to ensure consistency of fit. Results of fits shown in Supplemental Table 1.

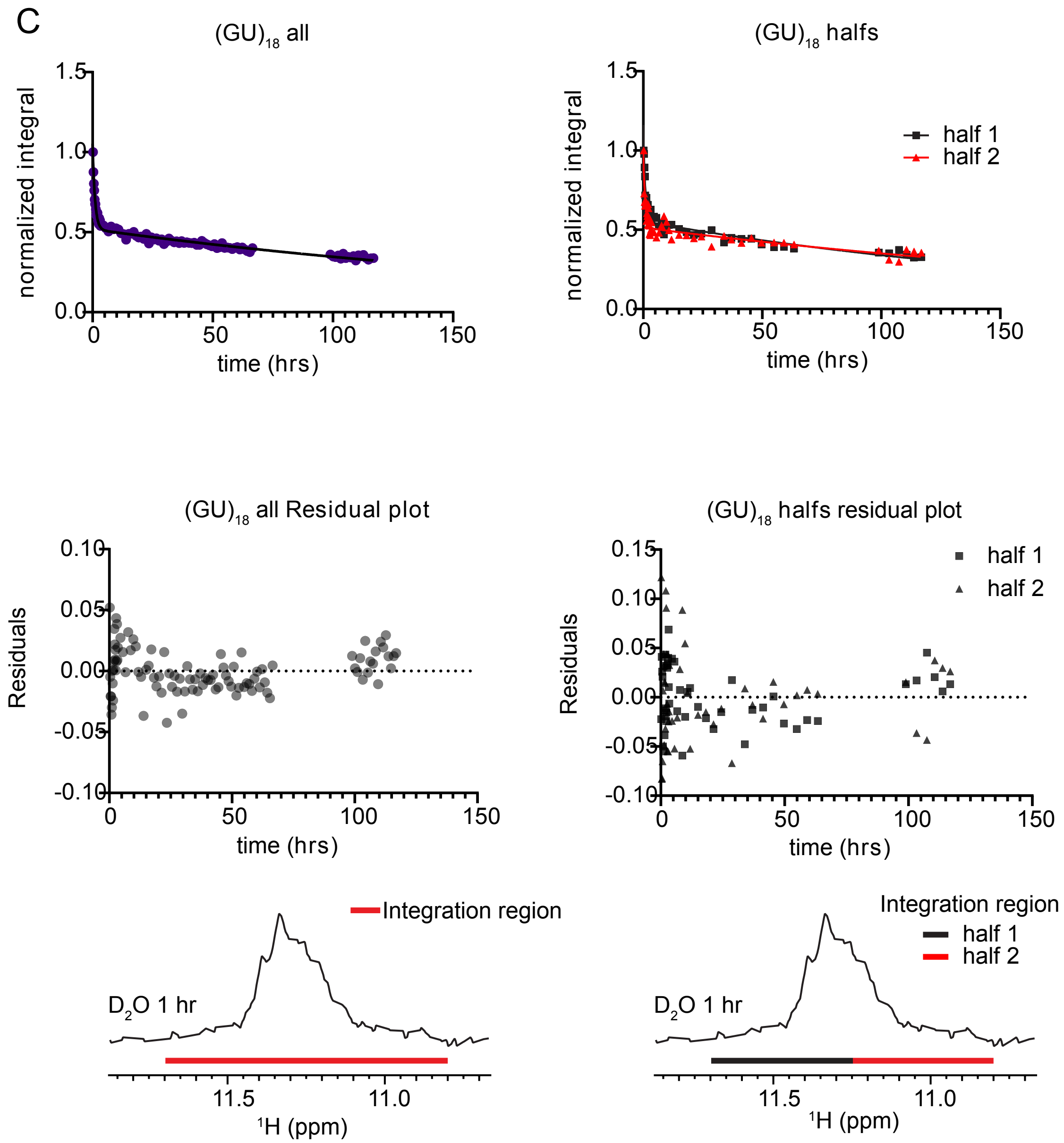


**Supplemental Figure 1C.** Global and regional fits of (A) $(GU)_{13}$, (B) $(GU)_{15}$ and (C) $(GU)_{18}$ HDX. From top to bottom: normalized integral (I(t)/I(0)) vs time plot fit to biexponential decay, residual plots, $^{1}$H NMR spectrum with integral regions shown. Left column shows entire imino region integrated together to account for broadening, right column shows split integral regions to ensure consistency of fit. Results of fits shown in Supplemental Table 1.

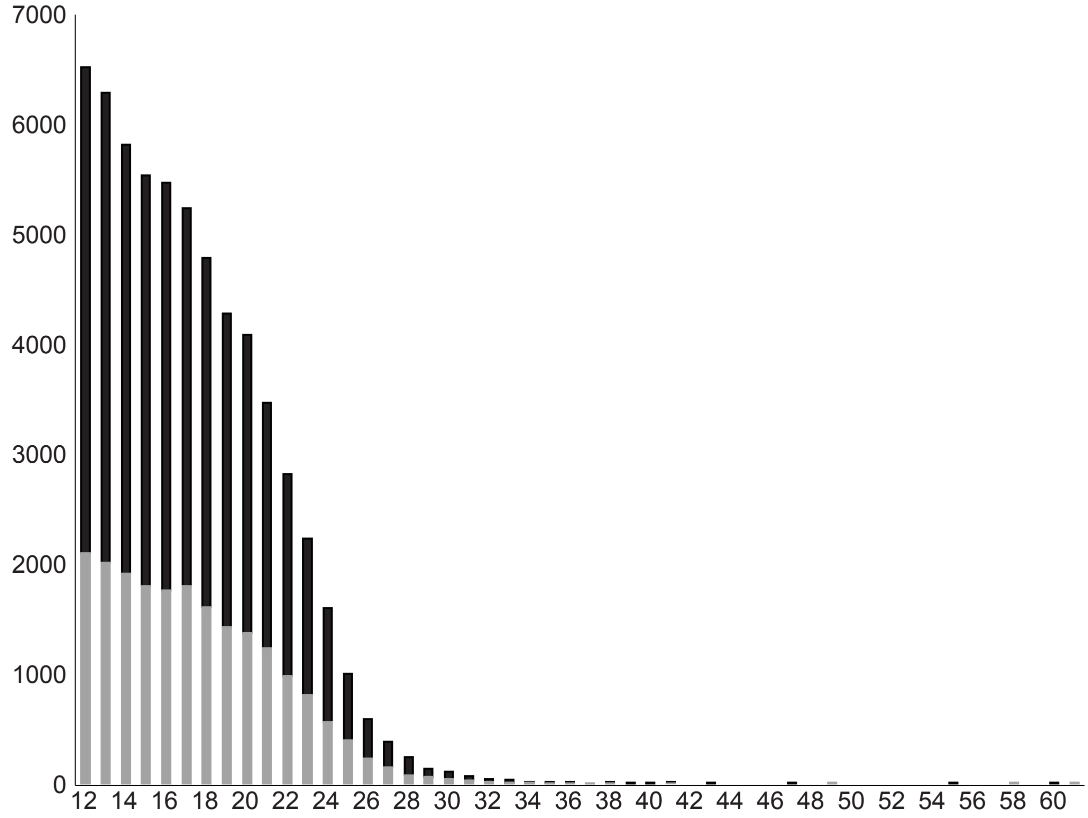


**Supplemental Figure 2.** Distribution of GT repeat lengths where n is the number of repeats. Graph includes ~60,000 GT repeat sequences containing >11 repeats, 22,000 of which are in annotated genes (grey). The average length is 17 repeats, with 4200 containing at least 24 repeats.

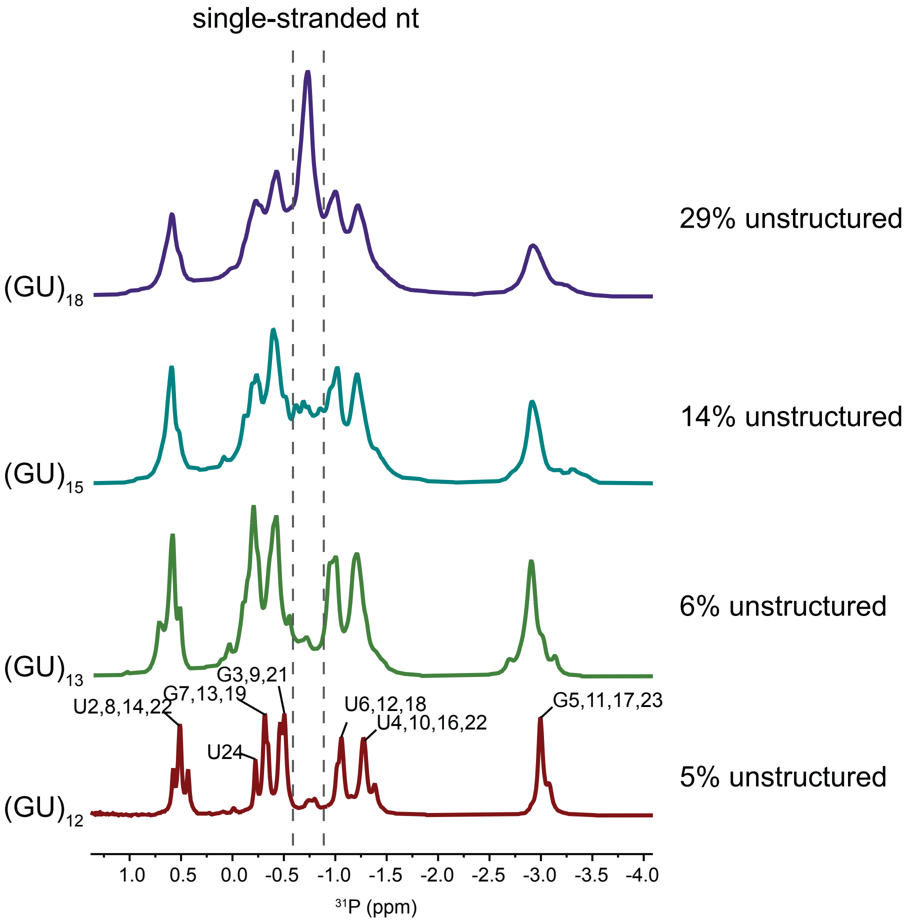


**Supplemental Figure 3.** 1D $^{31}P$ NMR of $(GU)_{12}$, $(GU)_{13}$, $(GU)_{15}$ and $(GU)_{18}$. Resonance assignments for $(GU)_{12}$ are shown. The peak at -0.75 ppm is attributed to unstructured nucleotides. Percent unstructured was calculated as the ratio of the integral of the unstructured peak vs. the integral of all the peaks.

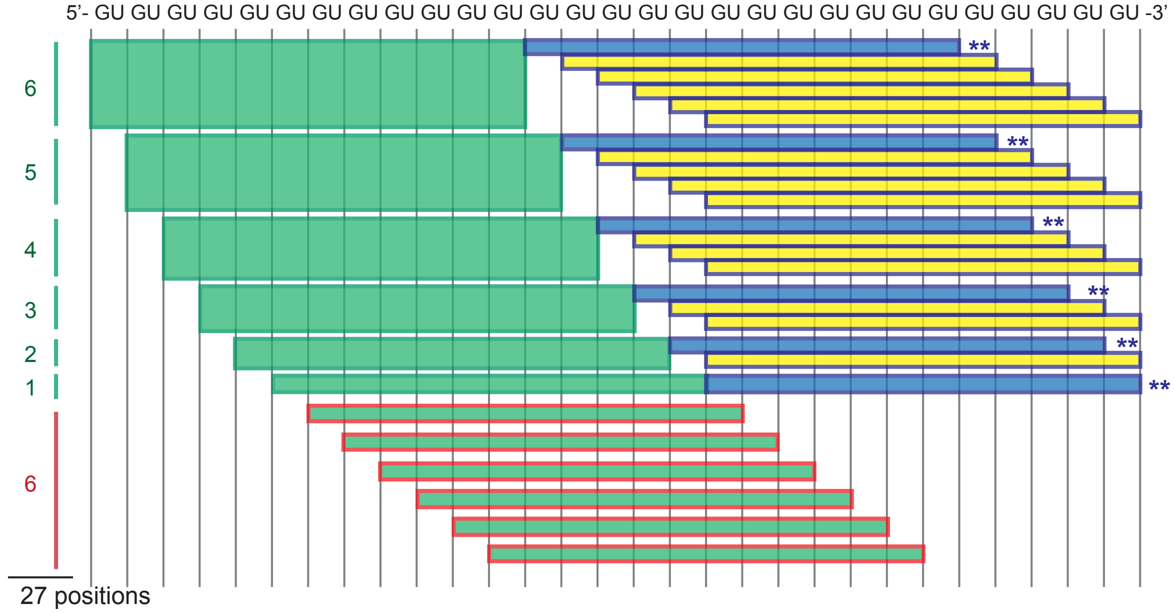


**Supplemental Figure 4** Diagram depicting all possible positions of pUG folds in $(GU)_{29}$ RNA. A pUG fold can occupy 27 alternative positions, 6 of which prohibit the folding of a second (green bars with red outlines). A stochastic model predicts 21/27 (77%) of $(GU)_{29}$ RNAs would have two pUG folds (either green and blue or green and yellow bars). Directly adjacent double pUG folds are shown as green and blue bars and are marked with 2 asterisks. Of all 27 possible combinations, 6 conformations, or 22%, include directly adjacent double pUGs.

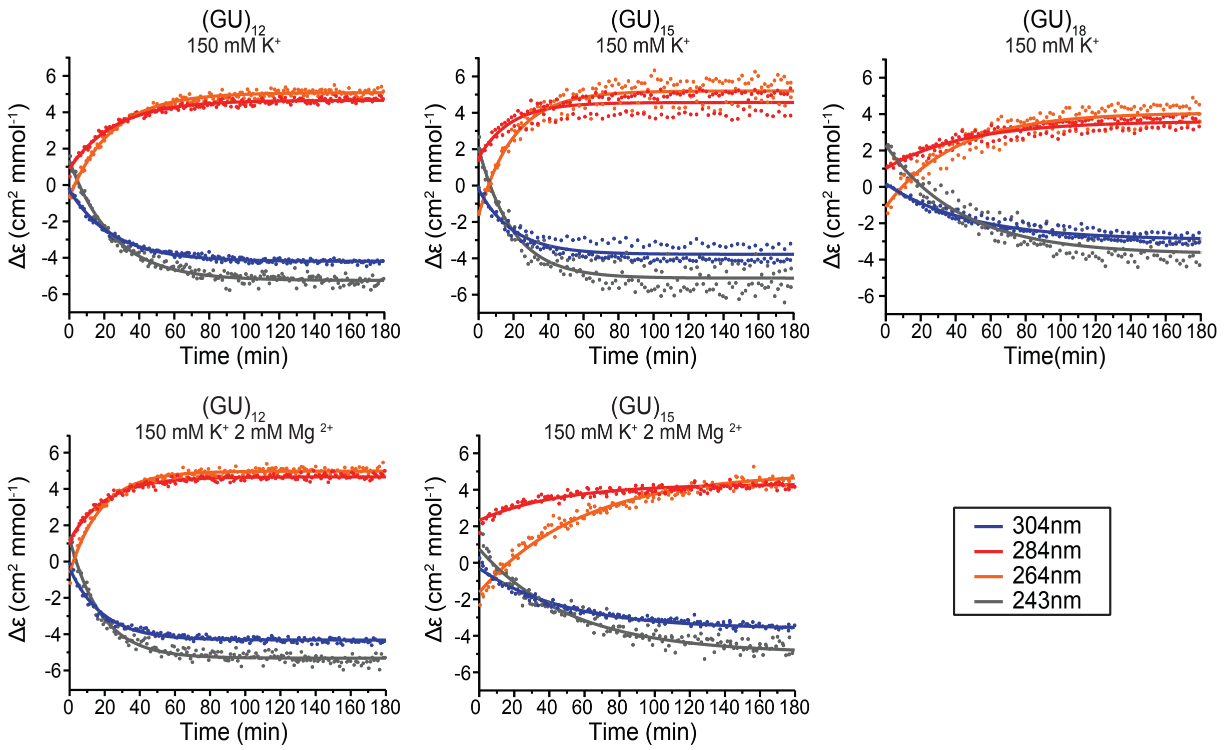


**Supplemental Figure 5.** CD monitored folding of $(GU)_{12}$, $(GU)_{15}$ and $(GU)_{18}$ initiated by the addition of 150 mM KCl containing buffer with or without 2 mM MgCl. The CD signal was measured at four wavelengths. Folding half-lives in $K^+$ are 17.5 ± 0.4 min for $(GU)_{12}$, 13.4 ± 0.5 min for $(GU)_{15}$ and 30.5 ± 2.1 for $(GU)_{18}$. Folding half lives in $Mg^{2+}$ are 13.1 ± 0.8 min for $(GU)_{12}$ and 35.6 ± 3.7 min for $(GU)_{15}$.

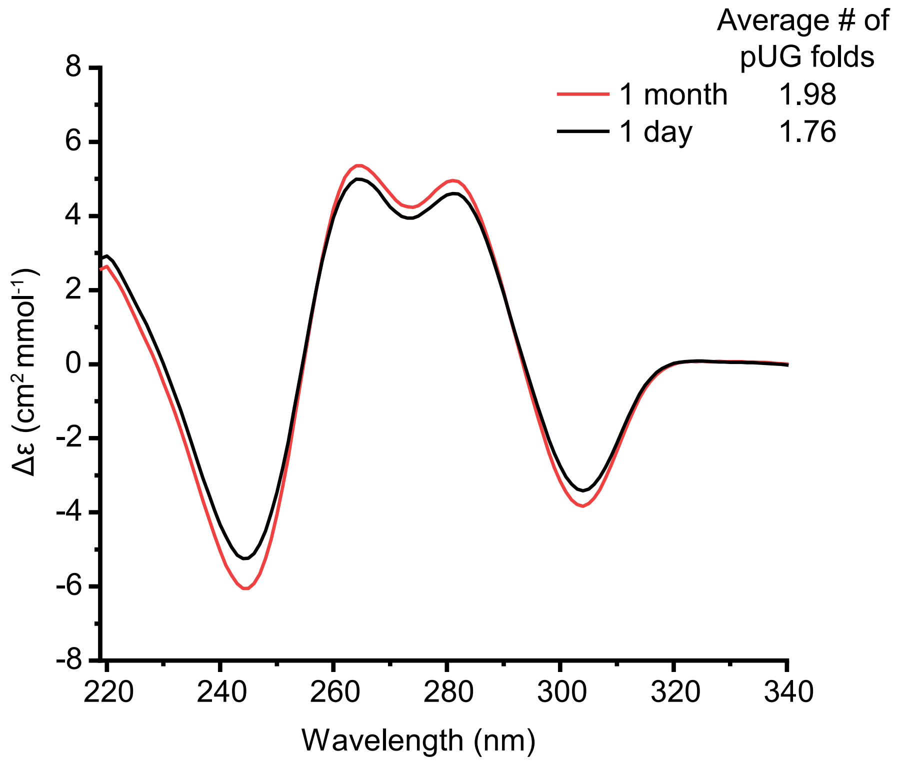


**Supplemental Figure 6.** Circular Dichroism spectra of $(GU)_{29}$ in 130 mM $K^+$ buffer following a heat and slow cooling folding protocol and incubation at room temperature for one day (black) or an additional month at 4 °C (red).

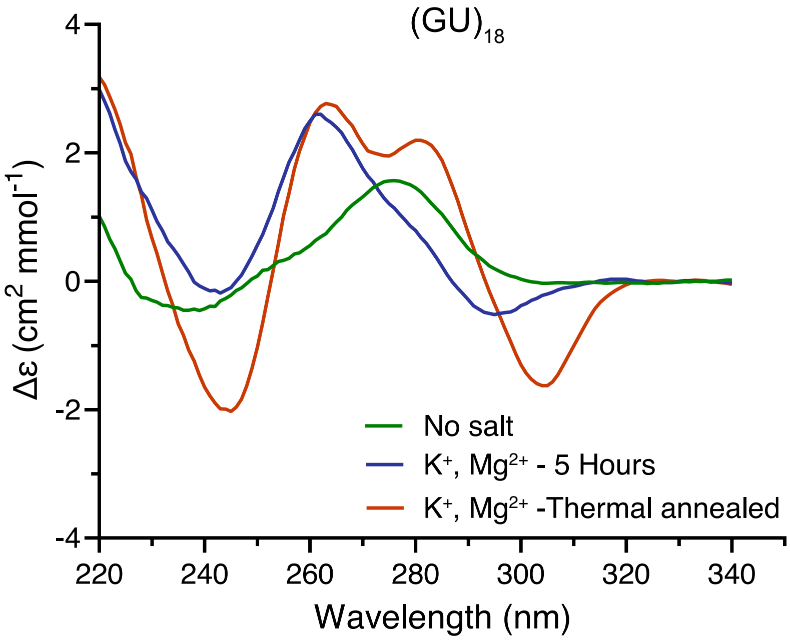


**Supplemental Figure 7.**
Circular Dichroism spectra of $(GU)_{18}$. Green, RNA in buffer with no added salt. Blue, RNA in $K^+$ and $Mg^{2+}$ buffer after 5 hours at 25°C. Red, RNA in $K^+$ and $Mg^{2+}$ buffer following a heat and slow cooling folding protocol.

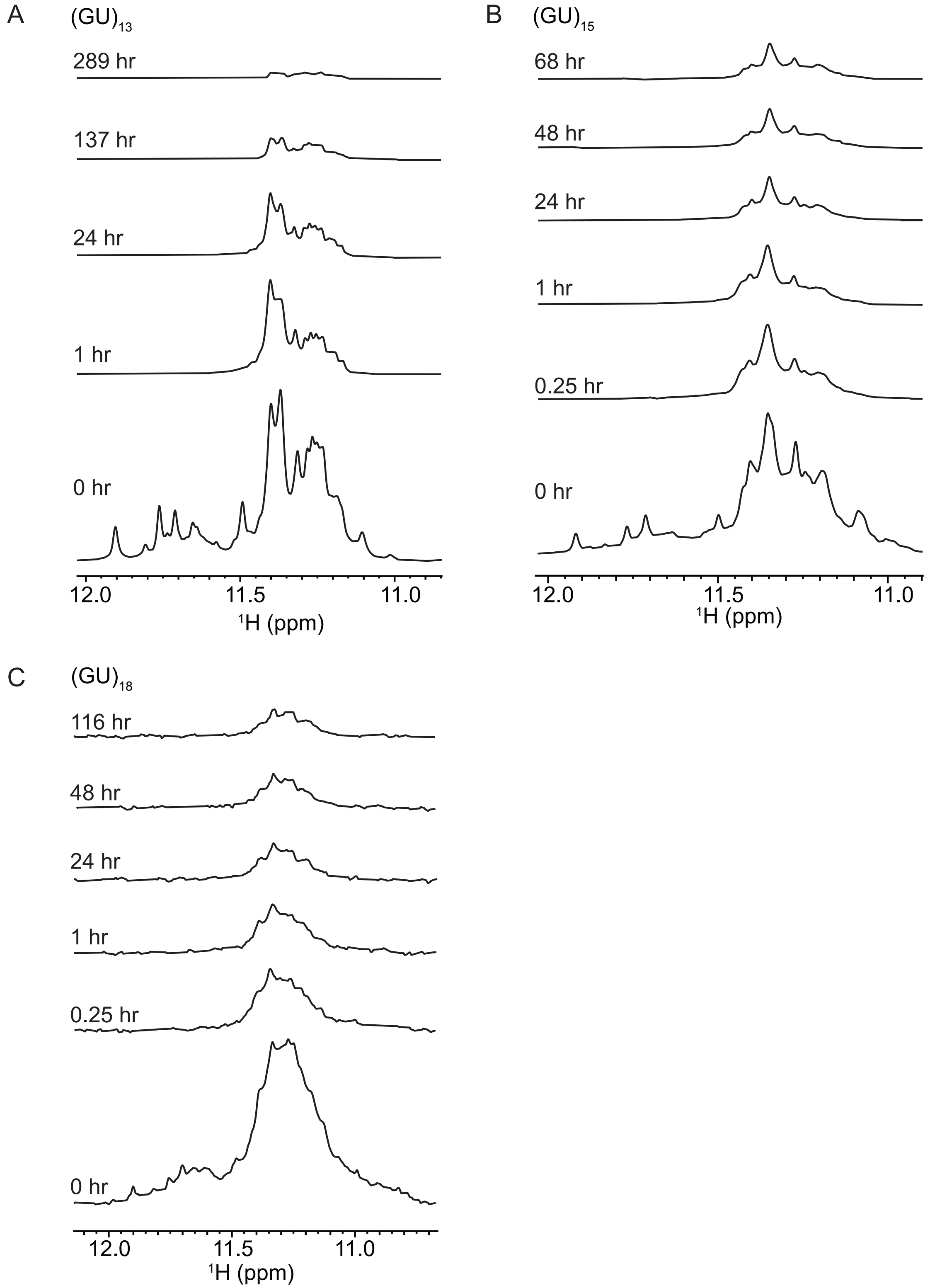


**Supplemental Figure 8.** $^{1}H$ NMR of (A) $(GU)_{13}$, (B) $(GU)_{15}$ and (C) $(GU)_{18}$ showing loss of imino proton signal at given time following transfer into $D_2O$.

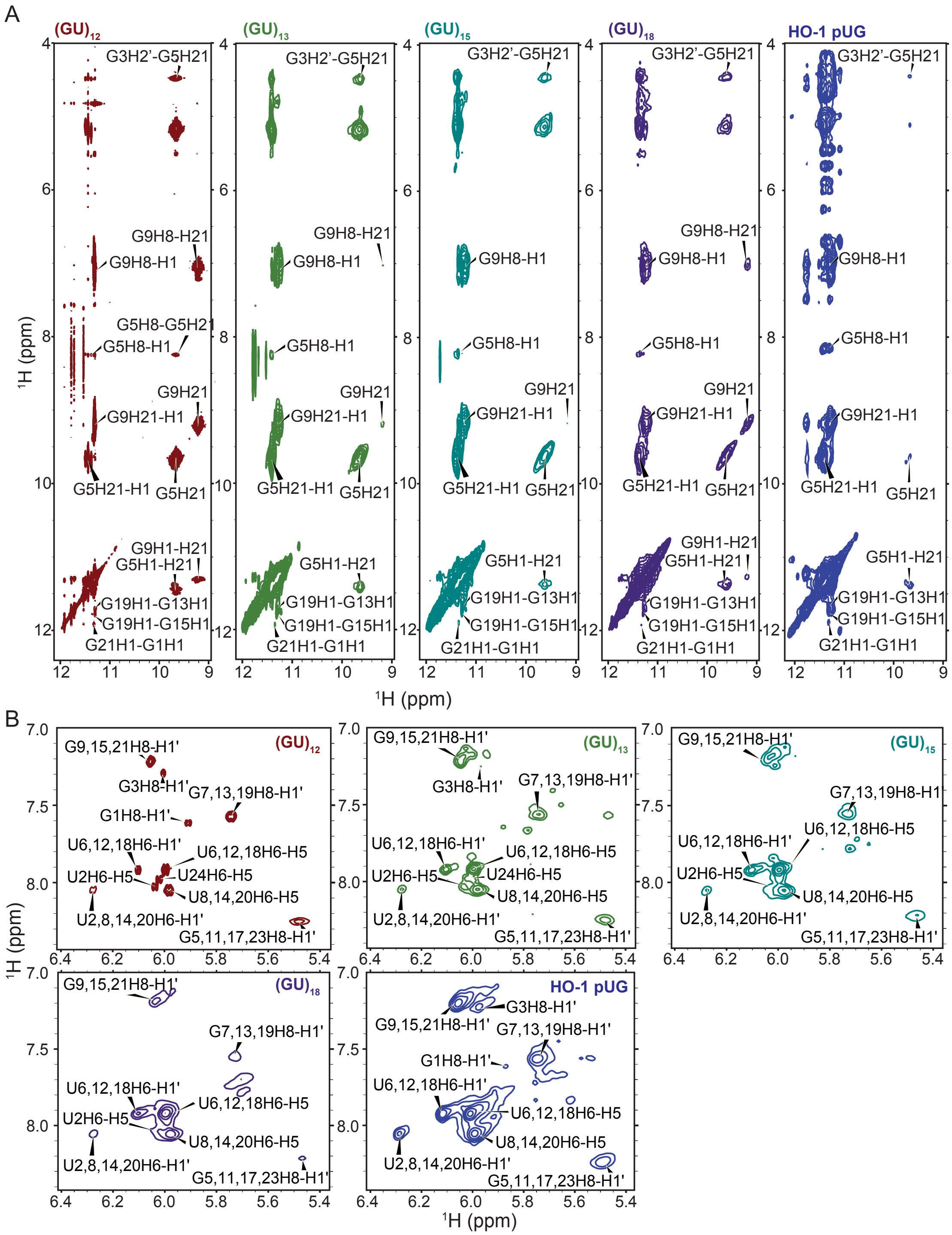


**Supplemental Figure 9.** 2D $^{1}H,^{1}H$ NOESY of $(GU)_{12}$, $(GU)_{13}$, $(GU)_{15}$, $(GU)_{18}$ and a 50 nt region of HO-1 pUG with the sequence $(GU)_{12}AU(GU)_{12}$. (A) NOEs of hydrogen-bonded imino and amino resonances and (B) aromatic and ribose NOEs. Peaks assignments are indicated.

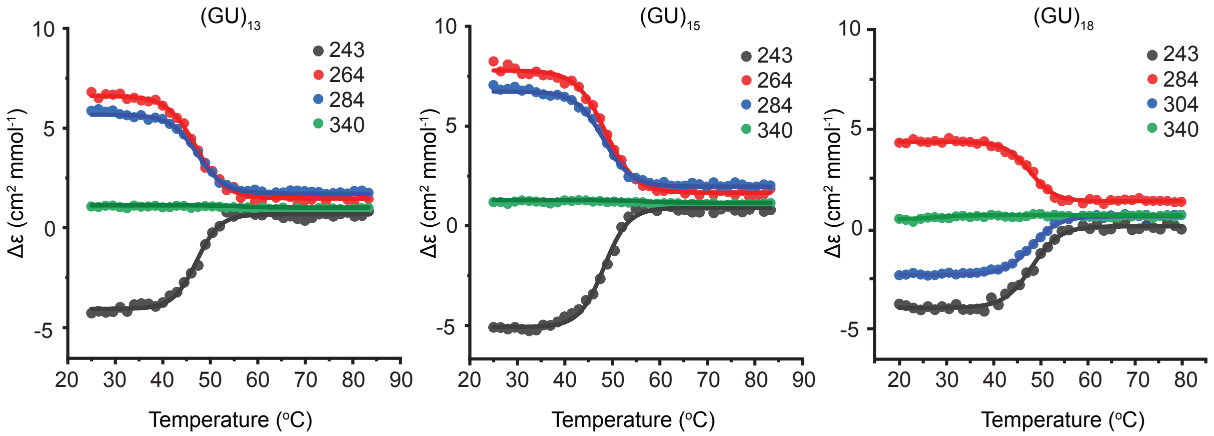


**Supplemental Figure 10.** CD monitored thermal denaturation of $(GU)_{13}$, $(GU)_{15}$ and $(GU)_{18}$ in 150 mM KCl. CD signal measured at three wavelengths plus a background control (340 nm) showing cooperative melting transitions. Melting temperatures are 48.3 ± 0.25 °C for $(GU)_{13}$, 47.1 ± 0.16 °C for $(GU)_{15}$ and 47.9 ± 0.18 for $(GU)_{18}$.

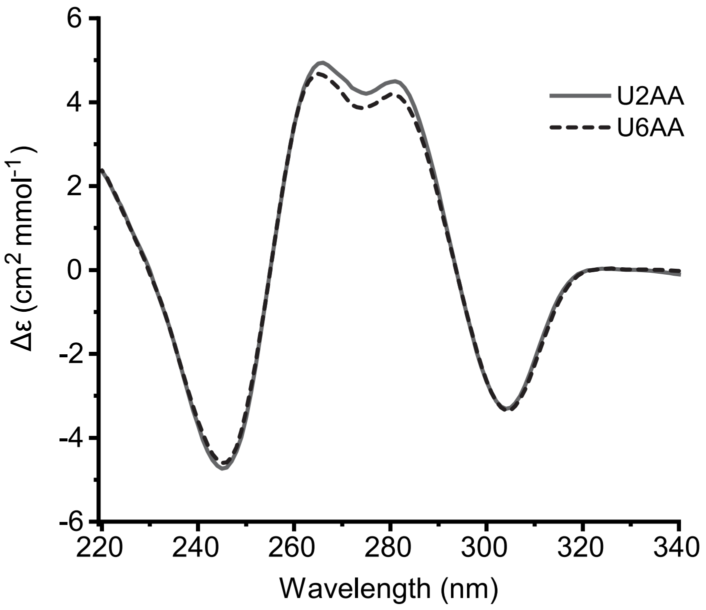


**Supplemental Figure 11.** Circular dichroism spectra of $(GU)_{12}$ containing AA insertions following the U2 and U6 position, $GUAA(GU)_{11}$ and $(GU)_3AA(GU)_9$ respectively.

**Supplemental Tables**

Supplemental Table 1: Divided fits of long pUG HDX

| | | I(0) | $k_{ex,fast}$ ($hr^{-1}$) | $k_{ex.slow}$ ($hr^{-1}$) | $t_{1/2,slow}$ (hr) | $t_{1/2,fast}$ (hr) | Ratio Fast (%) |
|---|---|---|---|---|---|---|---|
| $(GU)_{13}$ | Peak 1 | 1.09 ± 0.02 | 0.94 ± 0.06 | 0.009 ± 0.0002 | 76.4 ± 1.8 | 0.74 ± 0.05 | 42.4 ± 0.9 |
| | Peak 2 | 1.06 ± 0.07 | 0.97 ± 0.37 | 0.008 ± 0.0005 | 90.6 ± 6.2 | 0.72 ± 0.29 | 22.7 ± 5.1 |
| | Peak 3 | 1.10 ± 0.05 | 1.12 ± 0.16 | 0.007 ± 0.0004 | 97.0 ± 4.8 | 0.62 ± 0.09 | 41.6 ± 2.4 |
| | Peak 4 | 1.24 ± 0.06 | 2.19 ± 0.24 | 0.006 ± 0.0002 | 120.5 ± 3.9 | 0.32 ± 0.03 | 49.2 ± 2.4 |
| | Peak 5 | 1.31 ± 0.14 | 2.84 ± 0.61 | 0.005 ± 0.0003 | 126.9 ± 7.1 | 0.24 ± 0.05 | 48.1 ± 5.9 |
| $(GU)_{15}$ | half 1 | 1.06 ± 0.08 | 1.18 ± 0.27 | 0.006 ± 0.0014 | 122.3 ± 23.2 | 0.59 ± 0.13 | 53.1 ± 3.4 |
| | half 2 | 1.22 ± 0.15 | 2.01 ± 0.58 | 0.004 ± 0.0012 | 166.7 ± 36.7 | 0.35 ± 0.10 | 53.4 ± 6.0 |
| $(GU)_{18}$ | half 1 | 1.16 ± 0.08 | 1.20 ± 0.24 | 0.005 ± 0.0008 | 142.3 ± 20.5 | 0.58 ± 0.11 | 52.4 ± 3.4 |
| | half 2 | 0.99 ± 0.13 | 1.20 ± 0.45 | 0.004 ± 0.0013 | 182.2 ± 46.6 | 0.58 ± 0.25 | 48.0 ± 7.1 |
